# VmPFC supports persistence during goal pursuit through selective attention

**DOI:** 10.1101/2023.08.07.552276

**Authors:** Eleanor Holton, Jan Grohn, Harry Ward, Sanjay Manohar, Jill O’Reilly, Nils Kolling

**Affiliations:** Department of Experimental Psychology, University of Oxford, Oxford OX2 6GC, UK; Wellcome Centre for Integrative Neuroimaging (WIN), Department of Experimental Psychology, University of Oxford, Oxford OX2 6GC, UK; North Middlesex University Hospital, London, N18 1QX, UK; Nuffield Department of Clinical Neurosciences, University of Oxford, Oxford OX2 6GC, UK; Division of Clinical Neurology, John Radcliffe Hospital, Oxford University Hospitals Trust, Oxford OX2 6GC, UK; Centre for Functional MRI of the Brain (FMRIB), Wellcome Integrative Neuroimaging (WIN), Nuffield Department of Clinical Neurosciences, University of Oxford, Oxford OX2 6GC, UK; Université Claude Bernard Lyon 1, INSERM Stem Cell and Brain Research Institut U1208, NEF team, Bron, France

## Abstract

Striking the balance between persistence with a goal and flexibility in the face of better options is critical for effectively organizing behaviour across time. While people are often biased towards completing their current goal (e.g. ‘sunk cost’ biases), it is unclear how these biases occur at a mechanistic level, still allowing for some flexibility for goal abandonment. We propose that ventromedial prefrontal cortex (vmPFC) plays a critical role in orienting attention towards a current goal, prioritising goal completion but allowing for abandonment, particularly when the current goal fails. We developed a novel incremental goal pursuit task to study goal-directed attention and action in healthy individuals with functional magnetic resonance imaging (fMRI), and in an independent group of individuals with brain lesions. The task required participants to make sequential decisions between continuing to persist with a current goal (commitment), versus abandoning progress for a better alternative goal (flexibility). We show that individuals who persist more show greater goal-oriented attention outside the decision period. Increasing attentional capture by the current goal is also revealed in decision-making: people remain more likely to abandon from ‘frustration’ (collapse of value of the current goal) than from ‘temptation’ (attraction from valuable alternative goals). Strikingly, we find that our stable inter-individual metrics of persistence and goal-oriented attention were both predicted by baseline activity in vmPFC, tracking goal progress. We present converging evidence from an independent lesion patient study demonstrating the causal involvement of vmPFC in goal persistence: damage to the vmPFC reduces biases to over-persist with the current goal, leading to a performance benefit in our task.

## Introduction

In natural environments, many goals – whether it be pursuing prey, cooking dinner, or preparing an article for publication – are only obtained after persevering through a substantial period of unrewarded time and effort. In all these cases, optimal behaviour requires balancing commitment to the current goal against flexibility to abandon if the goal is no longer worth pursuing relative to alternatives. Psychiatry and neuroscience have tended to focus on *failures* of commitment during extended behaviours (LeHeron et al. 2019; Dalley & Robbins, 2017; Kouneiher et al. 2009). However, behavioural economics provides us with ample examples of people showing *too much* commitment to a goal, particularly after investing time or money (Arkes and Blumer, 1985; McAfee et al. 2010; Ronayne et al. 2021). Recently, it has been shown these ‘sunk-cost’ biases are not unique to humans, but exist in rodents too (Sweis et al. 2018).

Why might animals show biases towards over-persisting with a goal? When behaviour is structured by sequential goals, constant re-evaluation can be both expensive and distracting (Heckhausen & Gollwitzer, 1987; O’Reilly et al. 2020). In consequence, it has been proposed that distinct phases of ‘deliberation’ (evaluation of available options) and ‘implementation’ (committing cognitive resources to achieving the chosen goal) are present in both rodents and humans (Sweis et al. 2018; Ludwig et al., 2020; Li et al. 2019). However, a picture involving entirely discrete decision phases fails to explain how animals remain flexible to goal abandonment when the situation requires it. A plausible mechanism would allow for the agent both to preferentially allocate processing resources to goal completion, while retaining the necessary flexibility.

A candidate mechanism for such flexible focus on a goal is *selective attention*, specifically towards information about the chosen goal. Attentional selection need not be all-or-nothing, but can vary in strength as the need to exclude distractors varies (Lavie, 2005), thus allowing for flexibility. In ecological scenarios, we are faced with different reasons for abandoning a goal: progress might be too gradual or might reverse; alternatively other options might become significantly more attractive. These different forms of pressure give rise to different emotional responses: frustration (with the current goal) in the former cases (O’Reilly et al. 2020), and temptation (by alternative goals) in the latter. If selective attention to the chosen goal increases over the course of goal pursuit, this leads to a rather specific prediction about the interaction of ‘temptation’ and ‘frustration’ with increasing proximity to the goal: namely, sensitivity to the value of alternative goals (‘temptation’) should decrease more than sensitivity to the value of the chosen goal (‘frustration’). Our first aim was to test whether attention and decision-making showed these markers of increasing attentional orientation toward the current goal over the course of goal pursuit. To test this, we orthogonally vary the value of the current goal and the value of alternative goals at the decision, as well as continuously measure goal-oriented attention outside the decision period.

Our second aim was to investigate how goal commitment is achieved on a neural level. VmPFC has previously been shown to flexibly represent choice values according to the agent’s current goal (Grueschow et al. 2015, Rudorf & Hare, 2014, Castagnetti et al., 2021, Juechems et al. 2019, Trudel et al. 2021, Park et al. 2021), through the compression of task-irrelevant information (Mack et al. 2020). In addition to this body of research implicating vmPFC in task-specific cognitive maps, a separate line of research has identified a key role for baseline vmPFC activity in carrying state-specific information which biases subsequent choices in-line with a prior behavioural strategy (Lopez-Persem et al. 2016, Vinckier et al. 2018, Abitbol et al. 2015). While vmPFC represents attributes relevant to the current goal across extended time-scales (Korn & Bach, 2018), ACC has been shown to represent information about *alternative* goals and the value of shifting away from the current strategy (Blanchard and Hayden, 2014; Fouragnan et al., 2019; Hayden et al., 2011; Kolling et al., 2012, 2018; Tervo et al., 2021, Kaiser et al. 2021).

Using a novel task in combination with (i) computational modelling of behaviour, (ii) functional magnetic resonance imaging (fMRI) and (iii) behavioural analysis of patients with brain lesions, we investigated how goal commitment develops during goal pursuit. In our sequential choice task, participants advanced incrementally towards completing a chosen goal in the face of alternative goal offers. Participants showed a universal ‘goal commitment’ bias towards persisting with their current goal, even in circumstances when they would greatly benefit from abandoning it. We were able to measure several markers of selective attention to the current goal. First, as predicted by the attentional account, we found that decision-making reflected attentional goal capture: as participants approached goal completion, their decisions remained relatively more sensitive to the value of the current goal, than to the value of alternatives. Second, using a separate spatial working-memory task, we found that even outside the decision period, attention was increasingly captured by stimuli related to the current goal.

Using fMRI, we found that across participants, the degree to which baseline vmPFC tracked progress with the current goal predicted both attentional, and decision-based metrics of goal capture. To probe the causal role of this signal, we ran the same paradigm in an independent sample of patients with brain damage; indeed, damage to the same area of vmPFC identified in the fMRI study predicts lower commitment to the current goal.

## Results

Participants performed a “fishing net” task with the aim of filling as many nets with seafood as possible over the course of the study (Fig.1). Participants accumulated seafood “goods” over several trials, and only gained a reward when the net was full. On each trial, participants chose between offers for three types of goods (octopus, crab, or fish), where the quantity available for each good was shown by a green bar. Once selected, the offered quantity would be immediately added to the net. Importantly, only a single type of good could be collected in the net at once. This meant that if participants chose a different type of good to the type currently in their net, they would forfeit all their previously accumulated goods (‘abandonment choice’). Alternatively, participants could choose to continue with the current goal by choosing to collect the same good already in the net (‘persistence choice’; see Fig.1a for example).

**Figure 1.**
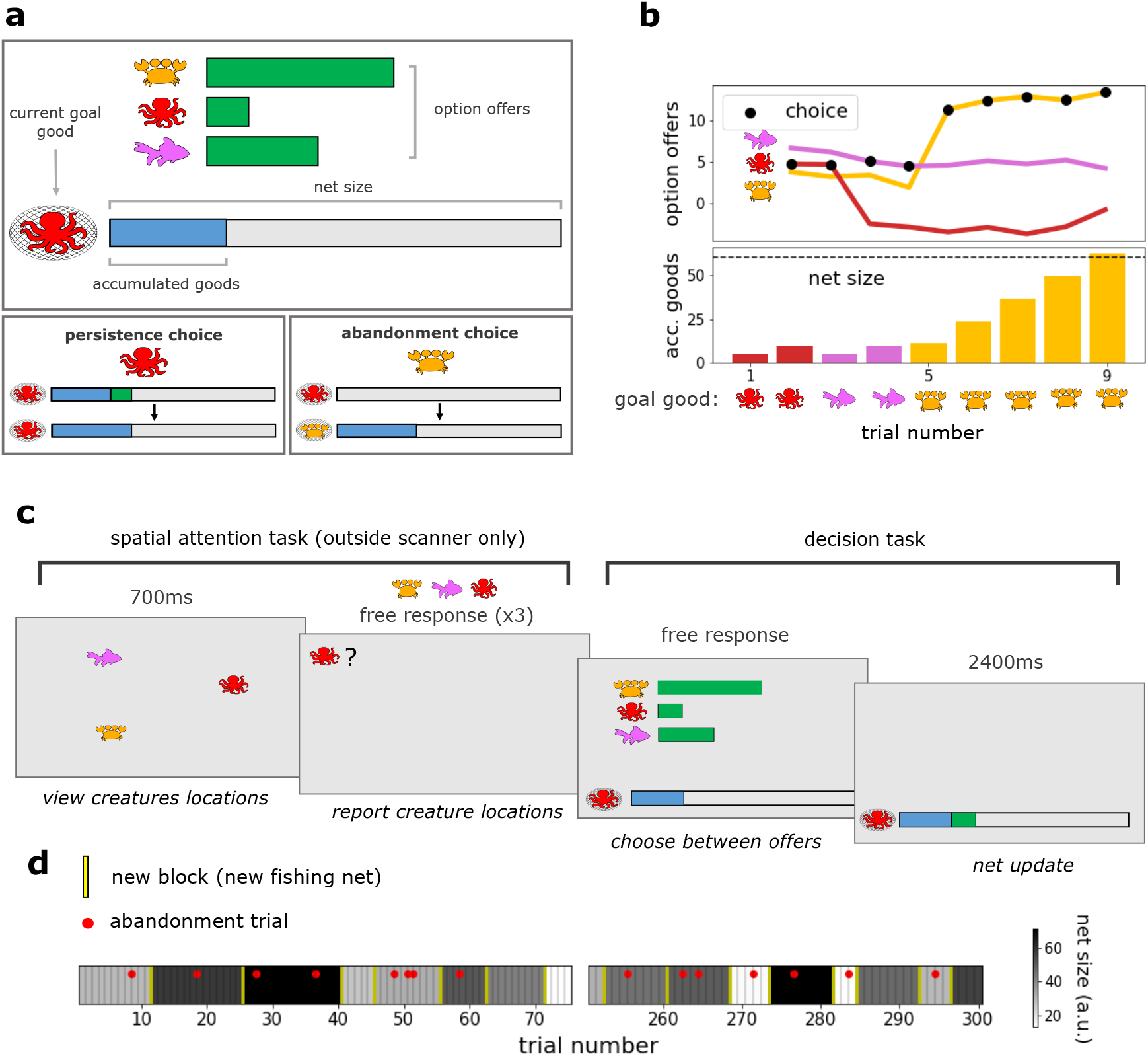
Experimental design. (a) Participants performed a “fishing net” task which involved accumulating goods over several trials, in order to be rewarded each time a fishing net was full. Top panel: on each trial, participants were offered the current available quantities of each type of good (octopus, crab, or fish) as bars on the screen. The length of each bar translated to the exact amount added to the net if that good were chosen, and the current contents of the net were shown in a separate bar at the bottom of the screen. Since only a single type of good could be accumulated in the net at once, switching between goods meant forfeiting the goods already in their net. Bottom left panel: If participants chose a good that matched the content of their current net, (‘persist’ trials), the offered quantity was added to the net contents. Bottom right panel: If participants chose a different good (’abandon’ trials), the net was first emptied of its accumulated contents before the offered quantity was added. (b) An example block where a participant switches twice. Top panel: Coloured lines depict the offers associated with each type of good across a block. Black dots depict the participants’ choice on each trial. During a block, the offers associated with each good varied across trials with independent random walks. In addition to this gradual variation, the offers could also jump to extreme high or low values, from where the random walk would continue. Bottom panel: Bars depict the accumulated goods in the net. Goods were accumulated until the net was full (shown by the dotted black line), triggering the end of the block and the delivery of a reward. Icons show the good currently accumulated in the net on every trial. Participants might switch because the goal good becomes worthless (example switch from octopus to fish) or because a different good becomes bountiful (example switch from fish to crab). (c) Task sequence. Outside the scanner, participants performed the same decision task with the addition of an interleaved spatial attention task performed on every trial. Participants viewed the three goods flash on the screen in random locations, and were then probed on the location of each good. Participants were made explicitly aware that performance in the spatial task had no impact on subsequent offers for the three goods. (d) Example experimental timeline. The task always ended after a pre-known number of trials (300 trials in the scanner, 100 trials in the post-scan session), incentivising participants to make strategic choices to maximise nets completed within the limited number of trials. Red dots indicate trials on which the example participant chose to switch between goods. Shade indicates the varying sizes of the nets. Yellow lines indicate when a net was completed. When a net was completed, this triggered the end of the block: a reward was delivered (one money bag per full net, regardless of net size or type of good), a new net was presented, and the quantities associated with each option were re-set.

While the quantities offered for each type of good drifted gradually from trial-to-trial (random gaussian walk with low variance), sometimes the quantity would drastically change for a given a good (10% chance of a large shift up or down in quantity, independent for each type of good; see Fig.1b for example offer trajectories across a block). If the quantity associated with the current goal good collapsed (causing ‘frustration’) or if an alternative good became much more bountiful (causing ‘temptation’), participants often benefitted from abandoning their progress and switching to an alternative good (Fig.1b).

### Attentional capture probe

Participants performed the decision task first inside the fMRI scanner, and then in a separate behavioural session outside the scanner. Outside the scanner, in addition to the main decision task, participants performed an interleaved spatial attention task before every trial, providing a separate measure of attentional capture by the current goal (Fig.1c, left). Participants viewed the stimuli associated with the three goods flash on the screen, and were then prompted to report the item locations with a mouse click (stimuli were probed in a random order). While the spatial attention task involved the same “seafood” stimuli, participants were explicitly told that memory performance would not impact subsequent offers in the decision task (See supplementary fig.1d,e for full illustration of the decision-only task in the scanner session, and the spatial variant of the task in the post-scan session).

### People are biased towards continuing with the current goal compared to an optimal model

Because of the need to commit to a good for many trials in order to realise the reward (delivered on the completion of a full net), a good decision is based not only on the current offer, but also the quantity already in the net and projections of future offers. To understand how participants made such projections, we constructed a series of models reflecting increasingly complex possible strategies (See Methods for details of models, model validation procedure, and model fitting procedure; see Supplementary fig.3a for graphic representation of models). Participants’ behaviour was best described by the most complex model we tested (“full task model”; Fig.2a). This model samples possible future trajectories for the option offers using the true generative procedure, and selects the option which is predicted to fill the net fastest when averaging across the sampled trajectories, providing an approximation of the optimal decision (monte-carlo sampling).

While general choice strategy was best described by the optimal model, people tended to over-persist with their current goal beyond the model’s predictions (Fig.2b; persistence biases were significantly greater than zero: *t*(29)=11.23, *p*<0.001 or Wilcoxon *T*=0.0, *n*=30, *p*<0.001). Persistence biases were quantified as the optimal model’s value of abandonment for which an individual is indifferent to abandoning (see green dots on Fig.2b). While by definition the optimal model is indifferent to abandonment at a value of zero, people tended to require a higher objective value of abandonment in order to actually abandon their current goal. This metric of persistence bias had excellent test-retest reliability within participants across sessions (Pearson’s *r*=0.70, *p*<0.001, see Supplementary fig.4d). Compared to the optimal model, persistence biases increased the more people progressed towards completion of the net (main effect of proportion of net completed on top of full-task model switch value: *X^2^*(1, *N*=30)=5.27, *p*=0.022; illustrated by binning in fig.2c; see Supplementary fig.3b,c,d for additional information about model confusion and analyses with goal progress).

### People who persist more also show greater attentional capture by the goal between decisions

We predicted that attentional and decision-making biases would be related during goal-pursuit. To measure attentional biases, we investigated how attention was distributed between stimuli associated with the current and alternative goals in a decision-free spatial attention task interleaved between decisions. Since the spatial attention task was not possible to perform using a button box inside the scanner, we investigated these attentional biases in a separate testing session conducted outside the scanner. In the post-scan session, trials of the spatial attention task were interleaved with new trials of the main decision task.

In the spatial attention task, participants were asked to report the location of briefly-flashed fish, octopus and crab symbols, using a mouse click. Indeed, participants were both more accurate and faster at reporting the location of the current goal stimulus compared to the alternative goal stimuli (Fig.2e; difference in accuracy for current goal vs alternative: *t*(29)=2.25, *p*=0.032 or Wilcoxon signed-rank *T*=130, *n*=30, *p*=0.035; difference in RT for current goal vs. alternative: *t*(29)=3.30, *p*=0.003, or Wilcoxon signed-rank T=85.0, *n*=30, *p*=0.002). This accuracy difference was primarily driven by progressive memory enhancement for the goal stimulus: spatial accuracy for the current goal stimulus increased with the number of trials participants had been pursuing the current goal (Fig.2f; effect of pursuit time on goal item accuracy: *t*(29)=-2.65, *p*=0.013, Wilcoxon *T*=121.0, *n*=30, *p*=0.021; there was no significant effect of pursuit time on accuracy for alternative stimuli: *t*(29)=-0.033, *p*=0.974, Wilcoxon *T*=205.0, *n*=30, *p*=0.584, *n.s*). In a direct comparison, there was a significant difference between slopes for the effect of goal pursuit on selected and alternative goal items (Fig.2f; *t*(29)=-2.37, *p*=0.024, Wilcoxon *T*=133.0, p=0.040). This effect occurred despite the fact that the task occurred outside the decision period, and that participants knew their performance on this interleaved task would not affect subsequent offers, suggesting a true attentional bias towards the chosen goal, that increases with goal commitment.

This metric of attentional goal capture directly predicted individual differences in persistence biases: people who showed more attentional capture by the current goal demonstrated higher persistence biases (Fig.2g; correlation between spatial bias and persistence bias. Note that this relationship holds even when attention-biases and decision-biases originate from separate behavioural testing sessions: Using persistence biases fit to data from scanner-only session: Spearman’s r=0.50, p=0.005; Using persistence biases from data aggregated across both scanner and post-scan sessions: Spearman’s r=0.53, p=0.003). This demonstrates that an individual’s tendency to over-persist with the current goal is related to their allocation of selective attention towards the current goal.

### Distinguishing two causes for goal abandonment: “temptation” vs “frustration”

How does progress towards a goal affect peoples’ sensitivity to the value of switching away to an alternative? We found that in general, people became less sensitive to the value of abandonment (defined as the projected value difference between staying with the current goal and switching to the best alternative goal) over the course of goal progress (i.e. interaction between abandonment value and proportion of net completed, on top of both main effects: *X^2^*(1, *N*=30)=42.43, *p*<0.001). We then asked whether this loss of sensitivity equally affected value associated with the *current* goal versus value associated with the *alternative* goals.

Pressure to abandon the current goal comes from two directions: an alternative good might become more attractive, pulling the agent towards the better option (‘temptation’) or the value of the goal good might collapse, pushing the agent away from the current goal (“frustration”; see Fig.1b for example). A rational agent should weigh these two forms of pressure equally when evaluating the options, since value is simply the estimated time in which the target can be completed with each option (i.e. already factoring in accumulated value; see supplementary fig.5c for simulation of optimal model on same task). Given that participants displayed increasing attentional capture by the current goal, we predicted that as a consequence, value associated with alternative goals would impact behaviour less than value associated with the current goal over the course of goal progress.

We found that people indeed showed an asymmetry in their use of these value sources which developed during goal pursuit. As an individual neared goal completion, abandonment was driven less by offers of highly attractive alternatives than by the current goal collapsing, compared to the normative model (Fig.2d). To test this, we predicted abandonment choices in a regression model using the interaction between goal progress and each source of value (alongside the main effects). Both sources of value impact behaviour less over the course of goal progress (interaction between alternative value and goal progress: *t*(29)=-7.97, *p*<0.001 or Wilcoxon signed-rank: *T*=10, *n*=30, *p*<0.001; interaction between current goal value and goal progress: *t*(29)=7.08, *p*<0.001 or Wilcoxon signed-rank: *T*=16, *n*=30, *p*<0.001). However, this loss of influence on behaviour affected alternative goal value more than current goal value (difference between alternative goal value*progress interaction terms and (sign-flipped) current goal value*progress interaction terms: *t*(29)=-3.39, p=0.002 or Wilcoxon signed-rank: *T*=77, *n*=30, *p*<0.001; visualised in Fig.2d by binning the data). In other words, over the course of goal pursuit, the impact of temptation from alternatives fades more rapidly than the impact of frustration with the current goal.

## fMRI results

### Inter-decision vmPFC activity tracks goal progress

Our behavioural analyses showed pervasive effects of goal pursuit on attention even outside the decision-making period. We reasoned that the brain regions involved in these attentional biases should similarly show goal-progress-related neural activity *that persisted outside the decision period*. We therefore conducted a whole-brain GLM analysis focussing on the inter-trial period. We modelled BOLD activity in the inter-trial period using regressors capturing an individual’s position in the goal (goal progress: proportion of target completed), the value of the current goal and the value of the best alternative in the previous trial (according to the full-task model), and the decision itself (binary abandonment vs. persist choice; see Supplementary fig.6a for correlation matrix and Methods section for full details of GLM). In addition, we controlled for decision-related activity by adding all of these regressors at decision time (time-locked to the onset of offers). The peak of activity tracking goal progress during the inter-trial period was in ventro-medial prefrontal cortex, vmPFC (Fig 3b).

**Figure 2.**
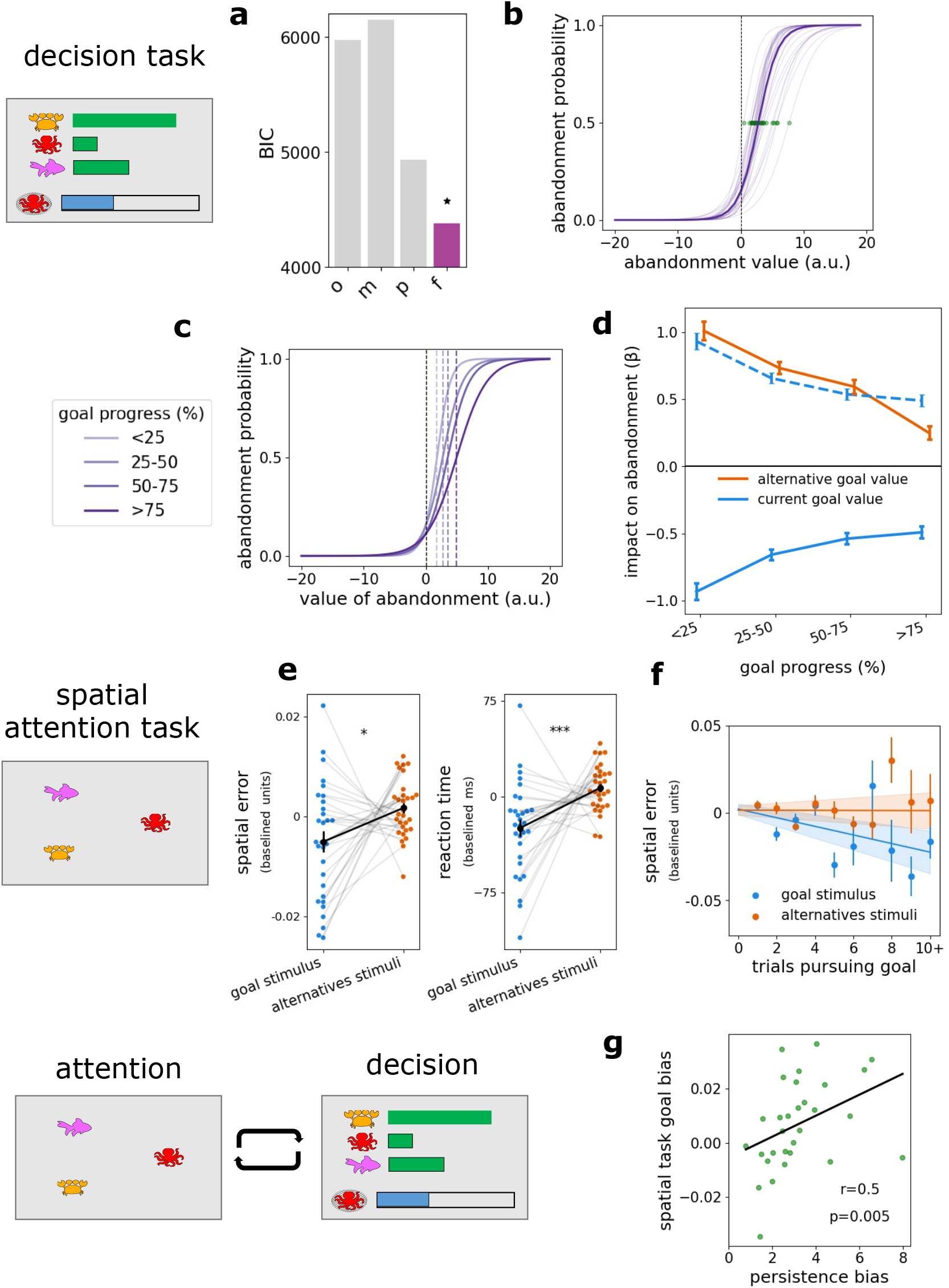
Behavioural results. ***Decision task*.** **(a)** Model fitting: The optimal model (full-task model; ‘f’) captured participant choices best (models compared using the Bayesian Information Criterion). See Methods for full details of the models and model-fitting procedure. **(b)** Persistence biases: When predicting peoples’ abandonment decisions using the optimal model value of abandonment, people showed a bias towards persisting. Bold line shows data fitted across all participants using a mixed effects model; Transparent lines show individual participant fits; Green dots show individual participant indifference points for abandonment which was used as an index of their ‘persistence bias’. Models fit to aggregate data across both sessions. **(c)** Across individuals, persistence biases increased as a function of goal progress (i.e. the percentage of the net which had been completed). Successive purple lines show sigmoid curves fitted using the same mixed-effects model procedure shown in (b), but this time binning the data by which quartile of the goal they were in, shown here for illustration. **(d)** Over the course of goal pursuit, the impact of temptation disappeared more than the impact of frustration on decisions to abandon. Blue and orange lines depict the beta weights associated with the optimal model value for the current goal and best alternative goal respectively, when predicting abandonment choices across goal pursuit. The dotted blue line is identical to the sign-flipped value of the solid blue line, shown here to illustrate the difference in slopes. Error bars depict standard error of the mean (SEM). ***Attention task*** **(e)** In the interleaved spatial task, both spatial memory error (left) and response reaction times (right) were lower for the stimulus associated with the current goal. Dots depict individual participants’ de-meaned error (left) or de-meaned reaction time (right), where blue shows the measure for the currently accumulated item, and orange shows the mean measure across the two alternative items. Error bars depict SEM. **(f)** As participants invested more trials in a particular goal, spatial error decreased for the current goal stimulus (but not for alternative stimuli). Dots show mean spatial error, binned by number of trials pursuing the current goal. Lines show the impact of number of trials pursuing the current goal, on spatial error, using the mean intercept and mean beta across regression models fit to each participant separately. Shaded regions depict SEM of the regression lines across participants. ***Relationship between decision and attention tasks*** **(g)** Individual tendencies to be more accurate at remembering the location of the goal stimulus than alternative stimuli in the spatial attention task (as depicted in Fig.2e, left) correlated with persistence biases in the behavioural task (as depicted by the green dots in Fig.2b). In other words, people who show greater attentional capture by the current goal tend to be more biased to persist with it. Here, persistence biases and attention biases come from data from separate testing sessions inside and outside the scanner respectively. Line depicts linear regression model fit.

**Figure 3.**
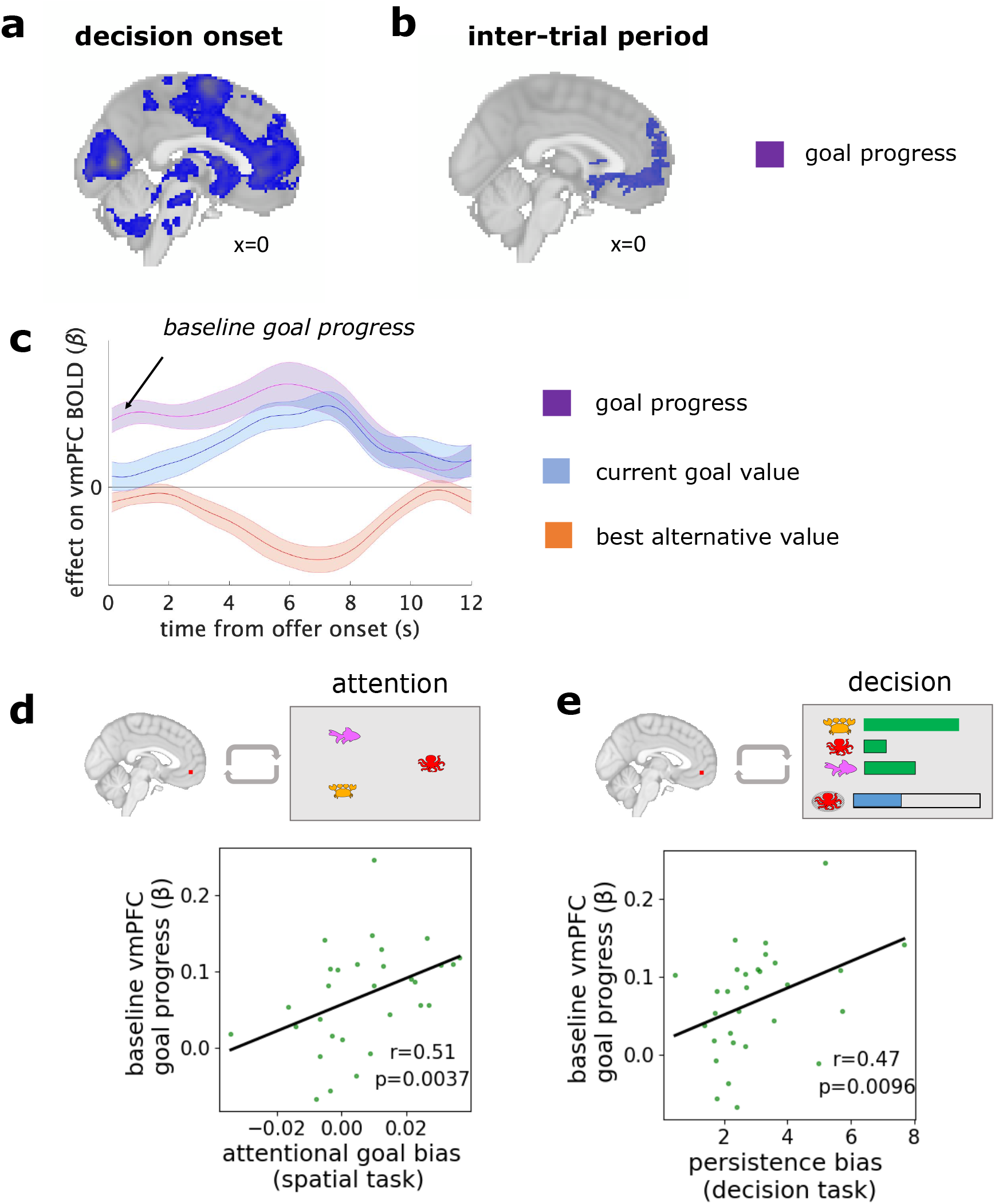
Goal-related baseline vmPFC activity correlates with individual differences in behaviour. **(a)** Cluster-corrected activity representing goal progress from the main analysis (time-locked to the onset of the decision period) **(b)** Same regressor as in (b) i.e. goal progress, but in an additional analysis time-locked to the inter-trial fixation cross (See *Whole-brain ITI analysis* in Methods for further detail). While there was widespread activity in in occipital and parietal areas representing goal progress at decision-onset (a), the majority of these areas were not involved during the ITI itself, where the highest peak was in vmPFC. **(c)** Time course of vmPFC activity in response to the onset of the offers, depicting the impact of the regressors of goal progress (purple), current goal value (blue), and best alternative value (orange) at decision time (beta weight on BOLD activity). Error bars show standard error of the mean across participants. Note that baseline activity tracks individual’s position in the goal (goal progress) before the decision is made. **(d)** Relationship between baseline tracking of goal progress in vmPFC and goal bias in spatial attention task (as in Fig.2e, left). Attentional goal bias refers to the accuracy advantage (in units on the screen) for reporting the location of the current goal item over reporting the location of the alternative goal items. Notably, the attention task measures come from an entirely separate session which took place outside the fMRI scanner. **(e)** Relationship between baseline tracking of goal progress in vmPFC and persistence biases in the decision task (as in Fig.2b, green dots; persistence biases fit across aggregate choice data from both sessions).

### Baseline vmPFC activity predicts the degree of goal-commitment across individuals

Previous studies have found that baseline vmPFC activity (before a decision) predicts biases or priors in decision-making (Vinckier et al. 2018, Abitbol et al. 2015, Lopez-Persem et al. 2016). As vmPFC maintains information about goal-pursuit between decisions, we hypothesised that the strength of this baseline vmPFC signal should predict the degree of commitment bias (unwillingness to switch goods) across individuals. We used a similar approach to previous studies by extracting baseline activity at the time of offer onset, on a trial-by-trial basis in our vmPFC region of interest (Vinckier et al. 2018). In all fMRI ROI analyses, we used regions of interest defined from orthogonal regressors related to value contrasts (see ‘ROI selection procedure’ in Methods).

We quantified the extent to which baseline activity in our vmPFC ROI varied with goal progress for each individual (quantified as the beta value within the ROI for goal progress, at choice onset). We found this baseline goal-related activity correlated with an individual’s overall persistence bias during the decision-making task (Spearman’s r(29)=0.47, p=0.010; Fig.3d).

If baseline vmPFC activity also reflects the degree to which the goal stimulus captures attention, we reasoned that it should correlate with the degree of attentional capture in the second, decision-free task. This was indeed the case – across participants the strength of the baseline goal-progress signal in vmPFC predicted greater accuracy for the current goal relative to alternative goals in the attentional task (Spearman’s r(29)=0.51, p=0.004; Fig.3e). This was particularly striking as the spatial decision-free task was carried out in a separate session outside the scanner.

Notably, both the reported relationships between neural activity and behavioural biases is specific to baseline activity in the vmPFC; baseline activity in other regions of interest and vmPFC activity in response to the decision itself are not predictive of behavioural biases (see Supplementary fig.8 for control comparisons).

### Influence of goal commitment on brain activity during the decision-process

As attention to the current and alternative goals varies with goal pursuit, we should expect to see changes in the representation of these goals at the time of the decision itself. We therefore asked how decision-related neural signals change as a function of goal pursuit. In particular, in behaviour we observed an intriguing asymmetry, namely that as goal commitment increased, sensitivity to alternative goal value (‘temptation’) was reduced more than sensitivity to the current goal value (‘frustration’). We therefore hypothesised that the neural representation of alternative value should change more with goal pursuit than the neural representation of the value of the currently pursued goal.

To identify brain regions involved in the decision process, we investigated neural activity at the time of the decision in a whole-brain analysis (regressors time-locked to the onset of the offers). This showed a much broader network of areas sensitive to goal-pursuit. Our whole-brain analyses revealed activity in a wide range of areas increased as an individual progressed towards completing the goal, including medial prefrontal cortex, striatum, and cingulate areas, as well as large regions of the occipital, and parietal cortices (‘goal progress’ regressor; Fig.3a). In addition, we found value-related activity consistent with previous findings related to brain networks involved in staying with a default versus switching to an alternative: both medial prefrontal cortex and striatum increased their activity as the value of persisting with the goal increased (value of current goal–value of best alternative; Fig.4a, blue). In contrast, ACC, presupplementary motor area (preSMA), bilateral dorsolateral prefrontal cortex (dlPFC), and bilateral insular, all showed the opposite profile: activity increased as the value of abandonment increased (value of best alternative–value of current goal; Fig.4a, orange), and activity was higher on trials where the participant chose to abandon the current goal (Supplementary fig.6b; See Supplementary tab.1 for activity peaks). Note that whilst a large number of neural areas tracked an individual’s position relative to goal completion during the decision, this activity persisted during the inter-trial period in only a subset of areas, focussed on vmPFC (‘goal progress’; Fig.3a,b) as previously described.

We subsequently selected regions-of-interest in three key value-sensitive areas for further analysis. VmPFC, ventral striatum, and dorsal anterior cingulate cortex (dACC) all showed strong value-related activity at decision time in our whole-brain analysis. This is consistent with previous literature showing dACC is involved in value-guided abandonment (Kolling et al. 2012, Fouragnan et al. 2019, Tervo et al. 2021), and ventral striatum is a centre of value-guided choice (Jocham et al. 2011), known to be sensitive to goal proximity (Howe et al. 2013), and with meaningful projections to vmPFC (Piray et al. 2016). Given the relevance of these areas for decision-making during goal pursuit, we created regions of interest at the peaks of activity in these areas from our whole-brain analysis. We then investigated whether the representations of chosen- and alternative value were stable or changed dynamically over the course of goal pursuit.

### Ventral striatum value signals reflect behavioural asymmetry in sensitivity to current versus alternative goals

We first tested for changes in activity reflecting the combined ‘abandonment value’ (the difference in value of the currently selected and best alternative goals) over goal pursuit, mirroring the initial behavioural analysis. We found that representations of the option value difference in the ventral striatum reduced as people progressed through the goal (interaction of value difference with goal progress: Wilcoxon signed-rank, *T*=126, *n*=30, *p*=0.028). There was no equivalent effect in vmPFC or ACC.

We then broke down ‘abandonment value’ into the value of the current goal and the value of the best alternative. Parallel with our behavioural results, we found an asymmetry between the impact of alternative and current goal value in the ventral striatum. Specifically, representations of alternative value disappeared in the ventral striatum over the course of goal pursuit, but activity continued to co-vary with the current goal value (Fig.4b, middle; orange line shows reduction in representations of alternative goal value, blue line shows stable representations of current goal value; interaction between best alternative value and goal progress: Wilcoxon signed rank, *T*=358, *n*=30, *p*=0.010; interaction between current goal value and goal progress: not significant; Wilcoxon signed-rank, *T*=167, *n*=30, *p*=0.178). This mirrored the behavioural phenomenon whereby people became relatively less sensitive to temptation by alternative goods, whilst maintaining sensitivity to the value of the chosen goal, over the course of goal pursuit. In contrast, there was no significant change in the representation of alternative value over goal pursuit in either vmPFC or ACC.

**Figure 4.**
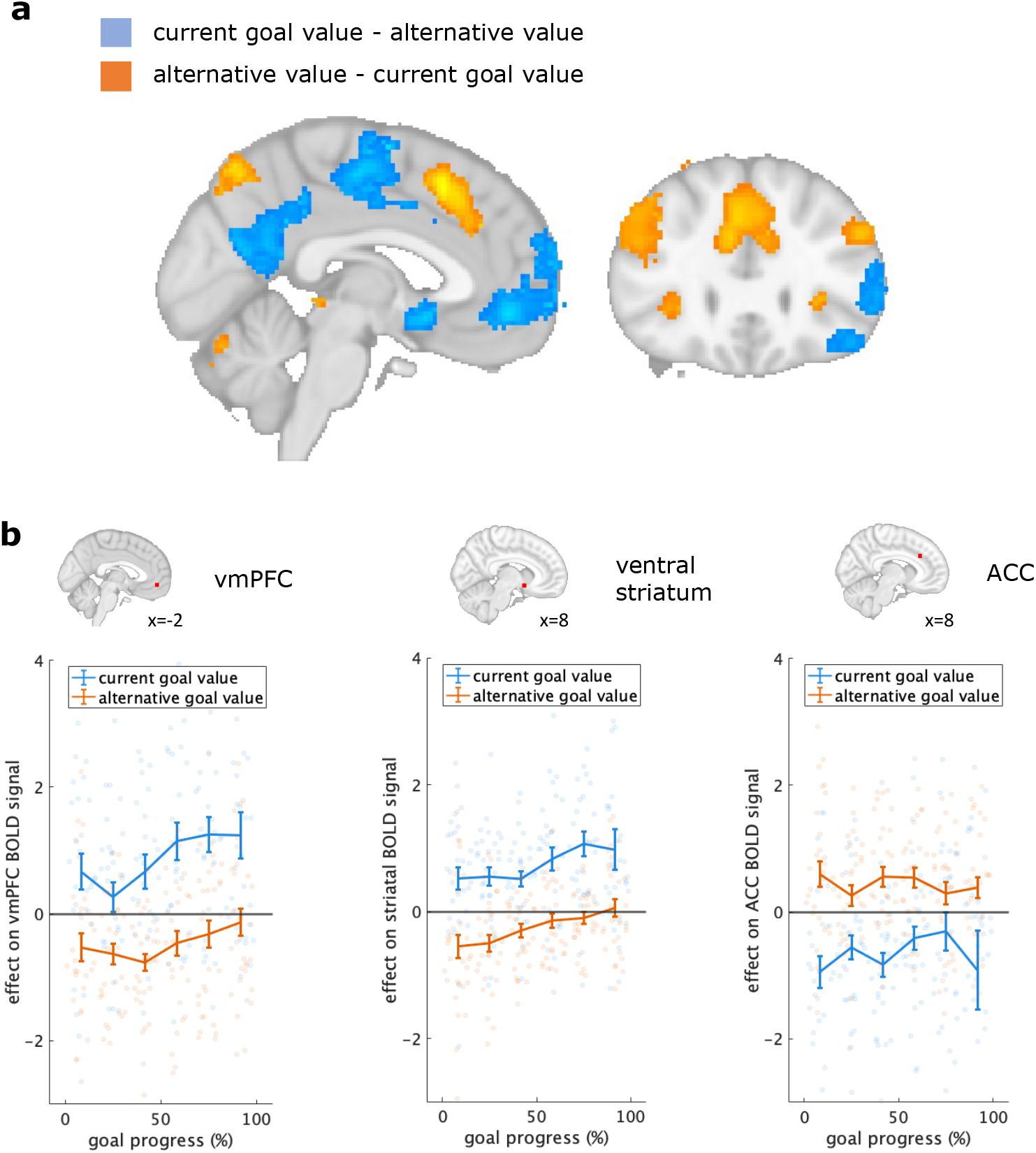
Neural activity related to the value of persistence and abandonment. **(a)** Results from the whole brain analysis showing cluster corrected peaks for the contrasts of current goal value - best alternative value (blue) and best alternative value - current value (orange). **(b)** Illustration of the modulation of value-related activity in our regions of interest over the course of goal pursuit. Here we show the effect of value on the BOLD signal (beta weight) against the proportion of the goal completed, binned for illustration. Error bars show standard error across participants, while dots show participant mean data points in each bin. Blue corresponds to the value of the current goal, orange corresponds to the value of alternative goals.

## Lesion Patient Study

### Damage to vmPFC reduces persistence bias for the current goal

Taken together, the behavioural and fMRI results suggest that vmPFC maintains attention to the chosen goal, leading to over-persistence or an unwillingness to switch goals. To test the causal nature of this association, we conducted an independent study using the same paradigm, with a sample of twenty-three participants with brain lesions in variable locations (see Fig.5a for map of lesion overlap across patients). We focussed on persistence bias, defined as the tendency to persist with the chosen goal beyond the point at which it would be optimal to switch, as the key behavioural marker of goal commitment.

We began by investigating whether damage to particular areas reduced persistence in the lesion patient group, independent from any priors from our fMRI study. We asked at what locations damage predicted a reduction in persistence bias by running a voxel-wise regression analysis using damage in each voxel (binary regressor) to predict persistence bias. Independently corroborating the findings of our fMRI study, the only region where damage predicted a reduction in persistence bias was in vmPFC (Fig.5b green cluster; cluster threshold *t*>2.3 (*p*<0.01, one-sided), cluster size=269 voxels, threshold cluster correction size=255 voxels, cluster peak=[0,42,-14], *t-*statistic at cluster peak=2.74, *n*=5 patients with damage within cluster).

We then asked how much the region identified in our lesion patient study aligned with the findings of our fMRI study. Our fMRI study had identified a subset of areas carrying signals relating to goal-pursuit even between decisions, focussed on vmPFC. We split all patients into two groups on the basis of whether they were damaged within a region of interest at the peak of this fMRI activity, found in vmPFC (region of interest centred on the peak of the activity tracking goal progress during the inter-trial interval in our fMRI study; shown in supplementary fig.9d). There were four lesion patients with damage to this region of interest, and this group had reduced persistence biases compared to both patients with damage elsewhere, and to age-matched healthy controls (Fig.5c, persistence biases among patients damaged within fMRI ROI: *n*=4, mean=2.33, std=2.31; persistence bias among other patients: *n*=19, mean=6.12, std=2.88; persistence bias among age-matched controls: *n*=27, mean=5.29, std=2.74; difference between vmPFC group and other patients: permutation test, difference in means=3.79, p=0.012, one-sided; difference between vmPFC patients and age-matched controls: permutation test, difference in means=2.97, p=0.023, one-sided). Strikingly, we found these four patients who had damage within the region pre-defined by our fMRI study corresponded to four (out of the five total) patients identified from our independent voxel-wise patient analysis. Therefore our fMRI study and lesion patient study independently converge to identify the same vmPFC region as being relevant for goal commitment.

Next, we ruled out the possibility that the vmPFC damaged group were simply performing worse in some general way, for example by making random choices or forgetting the goal. An important point to note is that, because participants in general over-persist, a reduction in persistence biases should actually lead to an *improvement* in task performance, *if* participants switch goals at points at which it is beneficial to do so (rather than making random switches due to, for example, task disengagement). This is exactly what we find: the five vmPFC-damaged patients identified in our voxel-wise analysis in fact perform significantly better than patients with damage elsewhere, and no worse than age-matched healthy controls (Fig.5d; performance is quantified as mean trials to fill a net, i.e. smaller values indicate goals are completed faster: average goal-completion time in vmPFC patients: *n*=5, mean=7.71, std=0.46; completion time in other patients: *n*=18, mean=8.47, std=0.70; completion time in age-matched controls: *n*=27, mean=8.03, std=0.73; difference between vmPFC group and other patients: permutation test (one-sided), difference in means=0.76, p=0.015; difference between vmPFC patients and age-matched controls: permutation test (one-sided), difference in means=0.32, p=0.190, *n.s.*).

Finally, we used further post-hoc analyses to verify that a) vmPFC patients were not responding more stochastically b) vmPFC patients were not using a different normative model to solve the task. We formally quantified stochasticity as inverse temperature, and found the vmPFC group showed no difference in inverse temperature compared to other patients or age-matched controls (Supplementary fig.9b; see recoverability of inverse temperature parameter in Supplementary fig.4c; inverse temperature in vmPFC patients: *n*=5, mean=0.57, std=0.04; inverse temperature in other patients: *n*=18, mean=0.51, std=0.22; inverse temperature in age-matched controls: *n*=27, mean=0.61, std=0.19; difference between vmPFC group and other patients: permutation test, difference in means=0.06, p=0.572, *n.s.*; difference between vmPFC patients and age-matched controls: permutation test, difference in means=0.04, p=0.633, *n.s.*). We also found that, like for the MRI participants, decisions for all three groups in our lesion study are best described by the full-task (optimal) model (Supplementary fig.9a), suggesting vmPFC patients were not using a simpler response strategy.

Taken together, these results suggest that patients with damage to this region of vmPFC are not simply using a different task strategy or responding more randomly, but instead are less *biased* toward over-persisting with a goal.

**Figure 5.**
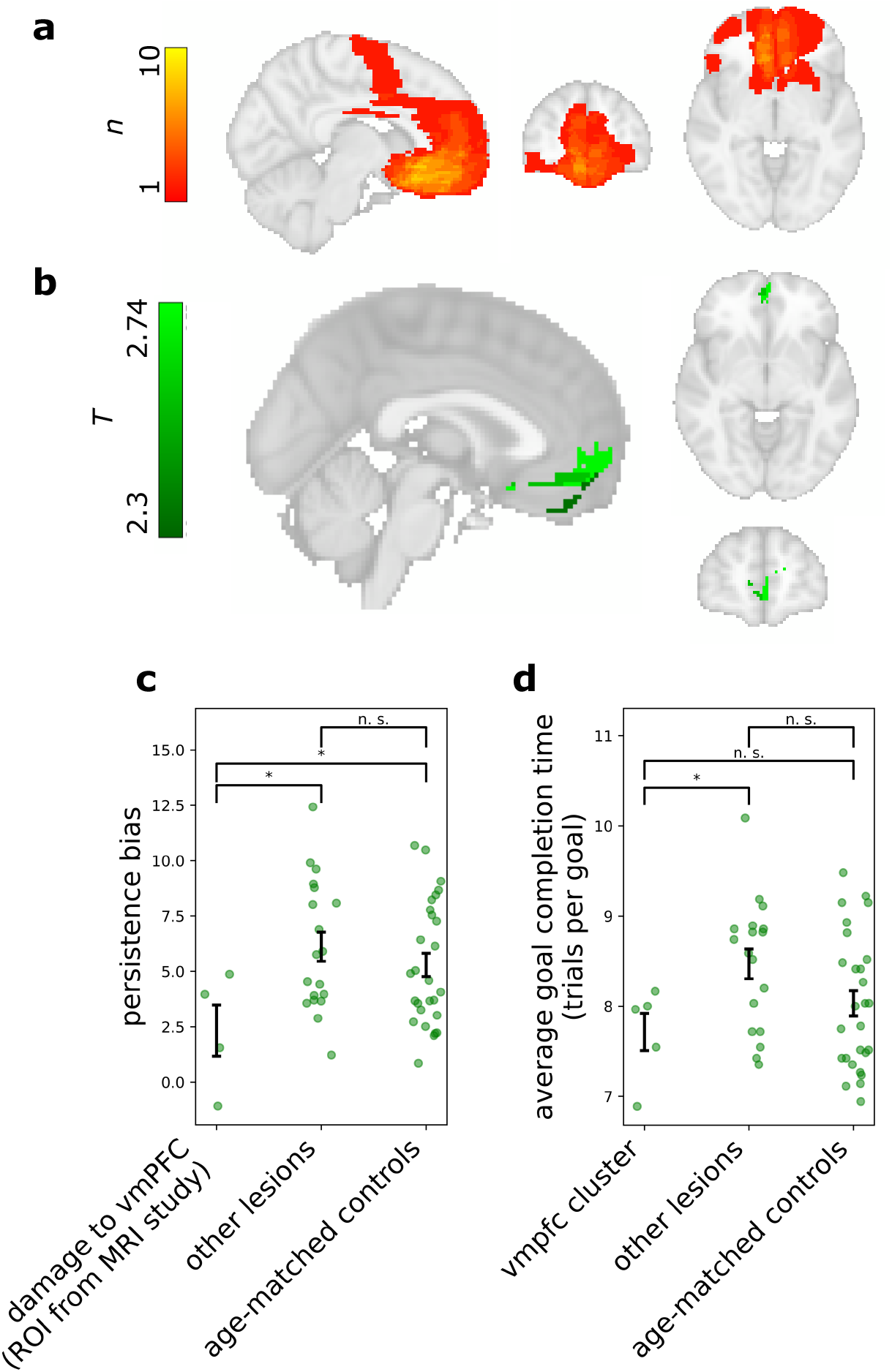
Lower persistence bias in patients with vmPFC lesions. **(a)** Lesion overlap maps of the twenty-three patients who took part in the study (maximum overlap in a voxel was 10 participants). **(b)** Green shows areas where damage predicts lower persistence biases in the patient study using a voxel-wise analysis approach. Here we show the map of t-statistics above threshold before cluster-correction for illustrative purposes. Only the vmPFC cluster survived whole-brain cluster correction as an area where damage leads to reduced persistence biases. The vmPFC region identified in the whole-brain voxel-wise analysis contains damaged voxels from *n*=5 patients (out of a total of *n*=23 patients who took part in the study). **(c)** After independently identifying the relationship between vmPFC damage and reduced persistence in our patient study (see b), we asked how much this result corresponded to the findings of our fMRI study. We split patients into two groups depending on whether they were damaged in the same voxels carrying activity related to goal pursuit between trials (region of interest centred on the peak of activity representing ‘goal progress’ at the inter-trial interval from the fMRI whole brain analysis; see Methods). Note that this ROI contained four patients, corresponding to four out of the five patients independently identified in (b). Patients with damage to this region were less persistent than both patients with lesions elsewhere and age-matched controls. Error bars show standard error of the mean in each group, green dots depict individual biases. **(d)** Post-hoc analysis comparing performance between patients with damage in the vmPFC cluster (from b, *n*=5) damage external to the cluster, and age-matched healthy controls. Performance is quantified as the average number of trials taken to complete a net, so lower scores correspond to faster goal completion. These results show vmPFC lesion patients are not simply responding more stochastically since their performance is enhanced compared to other patients, and no worse than age-matched controls. These results show the reduced bias to persist results in a performance enhancement relative to other lesion patients.

## Discussion

Many rewards are only obtained after a period of persistent effort. Therefore a key challenge for agents is to maintain a balance between commitment with the current goal and flexibility if it ceases to be worthwhile. The current study presents evidence that this challenge it met via an attentional mechanism. It is well known that people tend to over-persist with chosen goals (the ‘sunk cost’ fallacy). Rather than representing persistence biases as a (perhaps irrational) factor in the decision process itself, we argue that it is better understood in terms of a more pervasive attentional effect: Mechanisms of selective attention, mediated by vmPFC, prioritise processing of the current goal over alternative goals, resulting in reduced sensitivity to attractive alternatives (‘temptation’). This attentional bias is sustained in time and generalizes outside the decision context, as participants showed reduced sensitivity to sensory features of goal-irrelevant stimuli (such as their location in space), particularly as the goal state is neared.

We developed a pair of complementary tasks to measure how attentional and decision-making biases develop together during goal pursuit. In the decision-making task, commitment to a goal is required in order to realise rewards, but participants also need to remain sensitive to changes in the value of goods associated with the current and alternative goals. Participants tended to persist with goals longer than was optimal. As people progressed towards the goal, they became less sensitive to the value of alternative goods compared to the value of the goal good, suggesting an increasing focus of attention on the current goal as they neared goal completion. We further probed this attentional account by interleaving the decision-task with an unrelated and decision-free spatial working memory task. We found that participants were better able to recall the location of stimuli associated with the goal, and this tendency increased as they continued longer with the goal. Furthermore, there were stable individual differences in persistence with a goal, which were predicted by individuals’ sustained attentional capture by goal stimuli outside the decision period. Individuals who were more biased to persist with a goal showed higher goal-oriented selective attention, even when these metrics were captured in separate testing sessions and an unrelated, decision-free task.

We present multiple converging lines of evidence demonstrating vmPFC plays a key role in this process. First, our fMRI study found that vmPFC carries sustained goal-related information between decisions in our task, and baseline activity before the decision predicts the two independent behavioural metrics of goal capture: both an individual’s bias to persist with the current goal, and their bias to prioritise goal-related stimuli in attention. This was the case despite the fact that attention was measured during a separate task outside of the scanner. Second, we show that vmPFC is causally involved in goal commitment: patients with damage to the same region have reduced biases to persist with the current goal.

We find that across healthy individuals, baseline vmPFC activity (activity before a decision is made) predicts both decision and attention biases in our task. This builds on a growing area of study in both monkeys and humans finding that baseline vmPFC activity plays a role in influencing how options are processed and subsequently which choice is made (Vinckier et al. 2018, Abitbol et al. 2015, Lopez-Persem et al. 2016). Baseline vmPFC activity has been argued to bias upcoming choices in line with prior contextual factors, including both stable preferences (such as tastes for music or food genres; Lopez-Persem et al. 2016), and dynamic states (such as satiety or mood; Abitbol et al. 2015, Vinckier et al. 2018). Our results provide evidence that another dynamic state, namely goal pursuit, modulates behaviour through baseline vmPFC activity. We argue that our results also offer a possible mechanism for these effects: sustained vmPFC activity drives global changes in selective attention, affecting how options are processed and which decision is subsequently made.

In various contexts, medial prefrontal cortex has been shown to support the selection of goal-relevant information through the compression of irrelevant dimensions (Mack et al. 2020, Wilson et al. 2014, Mante et al. 2013), flexibly adapting to changes in the current goal (Grueschow et al. 2015, Rudorf & Hare, 2014, Castagnetti et al., 2021, Trudel et al. 2021), through the creation of task-specific ‘cognitive maps’ (Schuck et al. 2016, Park et al. 2021). Other studies have also linked vmPFC activity to visual attention, both responding to exogenous manipulations of attention (Lim et al. 2011; Hare et al. 2011), and in mediating visual attention (Wolf et al. 2014). Here we present results bringing together these distinct bodies of research, suggesting that the role vmPFC plays in selecting goal-relevant information can also be directly linked to visual attention. Our proposal that attention supports persistence by filtering goal-relevant information is consistent with reports that visual attention modulates goal-relevant information when de-coupled from value (Sepulveda et al. 2020, Glickman et al. 2018).

VmPFC could be varying with goal-relevant information without playing any causal role in the decision process. To test the causality of vmPFC activity in goal persistence we carried out an independent study using the same paradigm with twenty-three lesion patients. Through a voxel-wise analysis of damage in our patient sample, we identified a large vmPFC cluster in which damage predicted reduced persistence biases. The area identified in patients closely corresponded to the area involved in persistence among healthy individuals, providing striking evidence that vmPFC plays a causal role in goal commitment. Our results expand on previous reports that lesions to this area in both humans and primates interfere with the ability to prioritise the relevant decision variables, for example in cases when a distracting alternative is introduced (Noonan et al. 2010, Noonan et al. 2017), or an option has been de-valued (Reber et al. 2017).

While previous lesion studies have found this patient population to behave more stochastically (Noonan et al. 2017, Camille et al. 2011), notably, lower persistence biases among vmPFC lesion patients in our task cannot be explained purely by an increase in stochasticity. In fact we find patients with vmPFC damage performed better than other lesion patients and no worse than age-matched controls. In a goal pursuit context, healthy individuals may have a tendency to over-constrain the decision space by focussing only on the current goal and ignoring alternatives. In contrast, a lesion to this area of vmPFC may reduce selective attention to the goal, allowing alternatives to maintain their relevance throughout goal pursuit. We note that, while this is beneficial in our task, it is likely to be advantageous to constrain the task space in ecological goal pursuit settings, both in terms of optimal neural resource allocation (i.e. attending to goal implementation and avoiding cognitive switch costs), and in structuring behaviour over time.

Our results also reveal how neural value representations change dynamically across goal pursuit. We found that late in goal progress and compared to our normative model, people showed an asymmetry in how they weighed up value related to the current goal versus to alternative goals. Specifically, as goal completion was neared, participants were less sensitive to ‘temptation’ from attractive alternatives (compared to retaining sensitivity to ‘frustration’ should the value of the selected good suddenly fall). When the value of alternatives lost influence over behaviour, this was mirrored by a reduction in the representation of alternative value in the ventral striatum over goal pursuit. Striatal dopamine has been shown to ramp up during goal approach (Hamid et al. 2016; Howe et al. 2013), but also to trigger exploratory behaviours (Costa et al. 2014; Costa et al. 2019). It is possible these diverging functions could explain why we see representations of alternative options earlier in the goal, while striatal BOLD is saturated by representations relating to goal approach later on. Possibly, this asymmetry in striatal value representations could also reflect the increasing dominance of vmPFC during goal pursuit, reflected in striatal activity at a subsequent stage.

In contrast to the striatal effects, we found relatively sustained representations of alternative value throughout the goal in the ACC, supporting previous studies showing ACC drives flexibility. We found both ACC and dlPFC positively co-varied with the value of abandonment, as well as being more active when participants choose to abandon. This is consistent with previous work showing that activation in these areas, and in ACC in particular, represents the value of alternative options (Fouragnan et al. 2019), and is more active when an individual disengages from the present action (Stoll et al. 2016, Kaiser et al. 2021) or explores the environment (Tervo et al. 2021, Trudel et al. 2021). In fact, when people switch out of an exploitative state towards exploration, ACC activity predicts changes in task representation in vmPFC (Muller et al., 2019). While vmPFC represents the current goal and enables goal commitment, ACC is likely to underpin behavioural flexibility during goal pursuit by consistently tracking other options. The fact that people show increasing biases to persist rather than remain flexible could be explained by increasing dominance of regions such as vmPFC over ACC. We note that the vmPFC lesioned patients in this study made effective abandonment choices that allowed them to perform well in the task. While vmPFC contributes to persistence biases, it does not seem necessary for making good abandonment choices, which are likely to depend on areas such as ACC.

Our study suggests that goal pursuit involves the gradual shift of focus towards the current goal, rather than distinct modes of behaviour (“deliberation” and “implementation”) (Heckhausen & Gollwitzer, 1987, O’Reilly et al. 2020, Sweis et al. 2018). Selective attention provides a mechanism by which animals can prioritise goal completion while remaining sensitive to highly attractive alternatives, since attentional selection itself can be graded (Lavie, 2005). These persistence mechanisms which develop during goal pursuit and drive global changes in processing seem to be implemented through alterations in vmPFC activity across goal pursuit. While goal persistence may manifest in seemingly irrational tendencies to persist with a previous decision (as seen in classic “sunk-cost” effects), the ability to filter information to complete a chosen task is likely to be essential for adaptive behaviour in ecological settings.

## Acknowledgements

This research was funded in part by the Wellcome Trust (Grant number 222347/Z/21/Z to E.H.). For the purpose of Open Access, the author has applied a CC BY public copyright licence to any Author Accepted Manuscript version arising from this submission. This research was also funded in part by the Biotechnology and Biological Sciences Research Council (grant BB/ R010803/1 to N.K., https://bbsrc.ukri.org/) and European Union (ERC to N.K., FORAGINGCORTEX, project number 101076247). Views and opinions expressed are however those of the authors only and do not necessarily reflect those of the European Union or the European Research Council Executive Agency. Neither the European Union nor the granting authority can be held responsible for them. J.G. was funded by the Medical Research Council UK (MR/K501256/1 and MR/N013468/1) and St John’s College, Oxford. The patient study was also supported by an Oxford NIHR BRC, the McDonnell foundation, and MRC clinician scientist fellowship MR/P00878X to S.G.M. We thank the Wellcome Centre for Integrative Neuroimaging, Oxford, for Magnetic Resonance Research, supported by core funding from the Wellcome Trust (203139/Z/16/Z and 203139/A/16/Z).

## Author contributions

The following list of author contributions is based on the CRediT taxonomy (Brand et al. 2015). E.H., N.K., J.O. and J.G. contributed to conceptualisation and methodology. E.H. contributed to software, validation, formal analysis, investigation, and writing the original draft. N.K., J.G. and J.O. contributed to supervision and reviewing & editing the manuscript. H.W. contributed to investigation in the lesion patient study. S.G.M. contributed to supervision in the lesion patient study and reviewing and editing the manuscript.

## Data Availability

The data that supports our findings can be obtained from the corresponding author upon reasonable request.

## Code Availability

The code to replicate the analyses and figures shown in this paper can be obtained from the corresponding author upon reasonable request.

## Methods

### MRI study

#### Participants

A total of 30 participants (19 female; mean age 25 years, normal or corrected-to-normal vision) were recruited via email circulation on Oxford University mailing lists and social media. No participants were excluded from the recruited sample. Ethical approval for the study was obtained by the Oxford Central University Research Ethics Committee (Ref: R72921/RE001). All participants gave written informed consent before the experiment. Participants were paid £15/hour plus a performance-dependent bonus between £8-12.

#### Experimental procedure

The training, scan and post-scan task were all carried out in a single session lasting 2.5-3 hours total. Before the fMRI scan, participants were trained on the task for approximately twenty minutes. Participants practiced on three full example blocks (on average approx. 25 trials, dependent on performance) with the interleaved spatial attention task included, and one additional example block without the spatial attention task (scanner version). Comprehension questions were included at the end of training to ensure that participants had understood the task structure. Once this had been verified, participants entered the scanner and completed 300 trials of the decision task only (since the spatial task could not be performed with the button box inside the scanner) lasting 50-60 minutes (scanner session). Participants then completed the spatial variant of the task for an additional 100 trials outside of the scanner, lasting 20 minutes (post-scan session). Once the post-scan session was complete, participants filled out a short debrief questionnaire.

#### Primary decision task

Participants were told their aim was to fill as many nets with seafood as possible across the study, limited only by the number of choice trials in the study. The number of trials remaining in which the participants could continue to fill nets was shown in the top right corner of the screen throughout the study (Supplementary fig.1a). Above the indication of trials remaining was shown the number of points earned (nets completed so far), where each completed net was converted to a 25p bonus payment at the end of the study.

At the start of each block, participants were shown the size of the net to be filled as an empty grey bar at the bottom of the screen (Supplementary fig.1b). Blocks ended when a net was complete, and a point was won (Supplementary fig.1c). On each trial, participants were presented with three offers associated with the three sea creatures (always crab, octopus, and fish). Offers were shown as horizontal coloured bars on the screen next to their respective creature, where the size of the bar translated exactly to the quantity which would be added to the net if that creature was chosen. Offers were mostly positive (indicated by green bars), but could occasionally become negative (indicated by a red bar). If a negative offer was selected, the quantity of the bar would be subtracted from the net. Once a net was empty, nothing more could be lost so choosing a negative offer would lead to no change.

In the scanner, participants indicated which creature they wanted to accumulate using a button box where the first three buttons corresponded to the top, middle and bottom creatures on the screen. Outside the scanner, participants selected the creature by clicking with the mouse. Note that across all versions of the task, the horizontal order of the three creatures on the screen was randomised on every trial to avoid confounding persistence with motor perseverance. Once the creature was selected, the participant viewed the net being updated according to their choice.

#### Spatial variant

After completing the task for 300 trials inside the scanner, participants performed 100 trials of a spatial variant of the task outside the scanner. The spatial variant included an interleaved spatial attention task before every decision (Supplementary Fig.1e). Participants viewed the three creatures flash up simultaneously for 500ms in randomised locations across the screen.

Participants were then probed in a random order on the location of each creature. When the icon of each creature appeared in the top right corner of the screen, participants responded by using their mouse to click on the location at which they remembered it appearing. While it was not possible for participants to perform the spatial attention task inside the scanner (due to the impracticality of reporting three spatial locations on every trial with a button box), the scanner variant matched the basic structure of the spatial variant whereby participants passively viewed the three creatures flash on the screen at a random time during an inter-trial interval of between 2.5 to 8 seconds (Supplementary Fig.1d).

#### Schedules

##### Schedule generation procedure

For each block, the size of the net and the option offers differed. The net sizes were drawn from a uniform distribution (min=12, max=72). The initial values for the three options were drawn independently from a normal distribution at the start of each block (mean=6, σ^2^=1). From trial to trial, the offers for each option changed according to independent gaussian random walks (σ^2^=0.8). In addition, on each trial there was an independent probability of any option changing more drastically in its associated offer (with probability of 0.1 jumping up and 0.1 jumping down), corresponding to an option becoming significantly more ‘bountiful’ or ‘scarce’ for fishing opportunities. The jump function consisted of drawing a random value between 3 to 9 points higher or lower than the option’s starting offer, which corresponded to the new offer for that item. After a jump, the subsequent offers for that option would continue to change according to a random gaussian walk from the new starting location (see Fig.1b for example trajectories created using this procedure). In order to select pairs of net sizes and option offers for which completing the net was non-trivial yet feasible, we chose combinations where goals were completed in more than 3 trials and less than 15 trials when choice behaviour was simulated using the full-task model.

##### Schedule variants

To minimise schedule-specific artefacts, we generated 5 different schedules which each consisted of 45 blocks of 100 trials. A block ended when the net was filled so participants on average viewed only 7 trials per block before completing the net. For each MRI participant, separate schedules were randomly selected for the within-scanner and post-scanner sessions. In the lesion patient study, the same schedule was used across all individuals (including age-matched controls) due to the limited sample size for lesion patients. Each session ended after a pre-determined number of trials (300 in the MRI scanner, 100 in the post-scan session, and 250 for all participants in the patient study), so no participant was able to complete all 45 blocks of a schedule within the available experimental trials.

#### Behavioural models

We investigated participants’ choice strategy by fitting their behaviour to a set of possible models capturing different strategies. Four models with increasing complexity were tested as candidates for describing peoples’ subjective evaluation of the offers (see Supplementary fig.3a for a graphic depiction of the strategies):

1. *Offer-max model*: The agent chooses the largest offer on screen, regardless of the accumulated contents in the net. The values of the three items according to the model are equivalent to the current offers for each item.
2. *Myopic model*: The agent maximises accumulated value on the *current* trial. This means they will only switch if an alternative offer is greater than the combined contents of the net and the offer for the current goal item. The value of the goal item is equal to the accumulated value plus the goal item offer, while the value of the alternatives is simply equivalent to their current offers.
3. *Simple prospective model*: The agent calculates how much progress towards the goal each offer will entail, where progress is the proportion of the remaining unfilled net that will be completed after choice. Mathematically, the value of an option according to this model is the current offer for each option, divided by the quantity of net left to fill (when choosing that option). Intuitively, this model values each option based on the number of trials needed to fill the whole net, if the option values stay constant throughout.

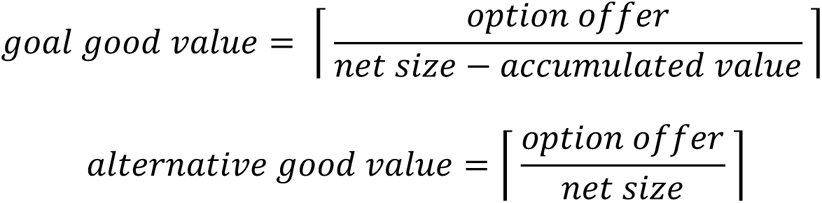 A central difficulty for a model which estimates value in this way is dealing with negative offers. Negative offers would *reverse* the respective values, meaning that implausibly, negative offers associated with the goal good are valued less than negative offers associated with alternative goods. To address this problem, we set the value of negative offers associated with alternatives to their raw (negative) offer, and the value of negative offers associated with the goal option to the proportion of progress they would be *losing* i.e. the offer divided by the accumulated value.
4. *Full task model*: This agent uses information about offer trajectories to simulate possible futures for the different candidate options, choosing the option which is forecasted to complete the net fastest. Specifically, it samples possible future trajectories for the three options and calculates each option’s value as the (negative) average number of trials until net completion across the iterations (if it were chosen on this trial). The same statistics used for creating the experimental offers were used when the model simulates the future trajectories of the options (procedure described in ‘Block Generation’). In other words, this model possesses task knowledge of how offers are likely to change over time, and leverages that to compute a better estimate of how long each option will take to fill the net.

#### Model fitting

Participant data was aggregated across the scanner session (300 trials) and post-scanner session (100 trials) before model fitting. In each case, the model value of switching was calculated as the model’s value for the current goal subtracted from the model’s value for the best alternative goal. To determine the best fitting normative model, we fit the following models to behaviour:

1. *SV_abandon_* = *β*_0_ + *β*_1_ * (*alternative value_offer-max_* – *goal value_offer-max_*)
2. *SV_abandon_* = *β*_0_ + *β*_1_ * (*alternative value_myopic_* – *goal value_myopic_*)
3. *SV_abandon_* = *β*_0_ + *β*_1_ * (*alternative value_prospective_* – *goal value_prospective_*)
4. *SV_abandon_* = *β*_0_ + *β*_1_ * (*alternative value_full-task_* – *goal value_full-task_*)

We fit these models in a mixed effects logistic regression analysis predicting abandonment choices, where intercept and slope were also modelled as random effects across participants. The Bayesian Information Criterion (BIC) was used to evaluate between models.

#### Model validation process

To validate the model selection procedure, we performed a model recovery analysis to confirm that the competing models were distinguishable within the empirical parameter range (Palmenteri et al. 2017). In addition to fitting the four basic normative models described above (see ‘Fitting Normative Model’), we also tested the recoverability of the basic models plus goal progress (See ‘Goal Progress’ section in Methods), which we found to have an additional impact on behaviour:

5. *SV_abandon_* = *β*_0_ + *β*_1 *_ (*alternative value_offer-max_* – *goal value_offer-max_*) + *β*_3_ ∗ *goal progress*
6. *SV_abandon_* = *β*_0_ + *β*_1 *_ (*alternative value_myopic_* – *goal value_myopic_*) + *β*_3_ ∗ *goal progress*
7. *SV_abandon_* = *β*_0_ + *β*_1_ * (*alternative value_prospective_* – *goal value_prospective_* + *β*_3_ ∗ *goal progress*
8. *SV_abandon_* = *β*_0_ + *β*_1_ * (*alternative value_full-task_* – *goal value_full-task_* + *β*_3_ ∗ *goal progress*

We used the empirical parameters from logistic regression models which were fit separately to each participant to simulate choices for each model. A soft-max function was then used to simulate choices from the subjective value:

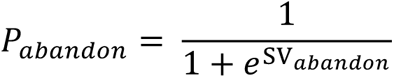

Subsequently, all models were fitted to all simulated datasets, and the empirical selection procedure applied to the simulated data (BIC comparison). To account for stochasticity resulting from the soft-max function, we repeated the simulation process 100 times for each of the 30 participants (resulting in 3000 simulated datasets per model). The averaged confusion matrix is displayed in Supplementary fig.3c, showing that each simulated model can be correctly identified during model recovery. Importantly, we find that in the empirical parameter range and across 100 repetitions, there are no cases of more simple models being confused for the empirically best-fitting model (full-task model).

#### Persistence bias

Once we established that the full-task model was the best fitting normative model of participant behaviour, we used this model to further investigate individual differences in persistence deviating from the model. We fit the logistic regression model defined in (4) of ‘fitting normative models’ to each participant separately. A participant’s indifference point (IP) is the model value of abandonment at which a participant is equally likely to persist or abandon (the ‘shift’ on the sigmoid function). Mathematically, this is equal to:

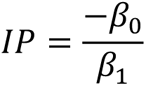

Where *β*_0_ and *β*_1_ refer to the intercept and slope respectively from the logistic regression predicting participant abandonment choices from the model value of abandonment.

Throughout subsequent analyses, the IP parameter fitted to each participant is referred to as their ‘persistence bias’, since all individuals had IP parameters above the value of 0 (i.e. they were biased to persist compared to the full-task model). See Supplementary fig.4a showing persistence biases can be accurately recovered after simulating behaviour within the empirical parameter range.

#### Persistence bias parameter recovery and test-retest reliability

We investigated the reliability of the persistence bias parameter, and the two sub-parameters from which persistence bias is derived (intercept and slope) using both simulated parameter recoveries and test-retest correlation across the two behavioural testing sessions (inside and outside the scanner). All three parameters show robust recovery in simulated data, as well as significant test-retest reliability in empirical data across the two behavioural sessions, as shown in Supplementary fig.4 (note that persistence biases have higher recoverability and test-retest reliability than either of its subcomponents on their own).

#### Goal progress

We define goal progress as the proportion of the current goal completed (i.e. current net contents / net size; Supplementary fig.3b). To quantify the additional impact of goal progress on peoples’ choices, we used chi-squared tests to determine whether each additional regressor improved our basic mixed-effects model across participants. For each model, the intercept and slopes for every regressor in the models were also included as random effects across participants.

1. *SV_abandon_* = *β*_0_ + *β*_1_ ∗ *V_abandon_*
2. *SV_abandon_* = *β*_0_ + *β*_1_ ∗ *V*_*abandon*_ + *β*_2_ ∗ *goal progress*
3. *SV_abandon_* = *β*_0_ + *β*_1_ ∗ *V*_*abandon*_ + *β*_2_ ∗ *goal progress* + *β*_2_ ∗ *goal progress* ∗ *V*_*abandon*_
4. *SV_abandon_* = *β*_0_ + *β*_1_ ∗ *goal value_full-task_* + *β*_2_ ∗ *alternative value_full-task_* + *β*_3_ ∗ *goal progress* + *β*_4_ ∗ *goal progress* ∗ *goal value_full-task_* + *β*_5_ ∗ *goal progress* ∗ *alternative value_full-task_*

After finding that value is used less over goal progress (model 3 above), we quantified the relative decrease in use of “current goal value” as opposed to “alternative goal value” by separating the value of abandonment into its two components. For each participant, we fit a logistic regression model which included the interaction between each source of value and goal progress (alongside all fixed effects; model 4 above). To determine whether there was a difference in how goal progress impacts the use of the best alternative versus the current goal, we used a Wilcoxon signed-rank test to determine whether there was a significant difference between the interaction coefficients for best-alt*goal-progress and for (sign-flipped) current-goal*goal-progress across individuals. We tested against the *sign-flipped* coefficients for current-goal*goal-progress because the value of the current goal and the value of the best alternative have opposing impact on the likelihood of switching (see Fig.2d).

#### Spatial task analyses

The spatial task results come from a separate behavioural testing session after the fMRI session, where participants performed the same decision task with the addition of an interleaved spatial attention task before making each decision (the ‘spatial variant’ described above). We used this task to measure the relative distribution of attention between stimuli associated with the current goal, and stimuli associated with alternative goals, across goal pursuit. We quantified spatial error as the Euclidian distance between the location of the participant’s click and the true location at which the stimulus appeared, in normalised screen units. We quantified reaction times (RT) as the time in milliseconds (*ms*) between when a stimulus was probed (appearing in the top left corner of the screen), and when the participant indicated their response. To remove differences in baseline error and reaction time between participants, we subtracted each participant’s mean error and mean reaction time from all their responses.

We then categorised responses according to whether the probed stimulus was the current goal good or one of the alternatives. We excluded the first trial of every block from analyses, where no goods had yet been accumulated. To quantify goal biases within participants, we took the difference between their mean behavioural response measure (normalised error and normalised RT) for the current goal stimulus, and their mean for the two alternative stimuli (Fig.2e).

We then investigated whether the spatial error bias developed as a function of goal pursuit (Fig.2f). We fit two linear models for each participant predicting (a) current-goal stimulus error and (b) alternative stimuli error using the number of trials participants had been pursuing the goal:

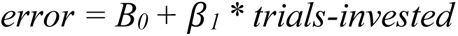

Using one-sample Wilcoxon signed-rank tests we investigated whether the *β_1_* coefficients across participants were significantly different to zero for either the current-goal stimulus or the alternative goal stimuli. We then used the Wilcoxon signed-rank test to determine whether the coefficients were significantly different to each other.

Finally, we investigated whether goal biases in the spatial task were related to persistence biases in the decision task. We tested for a relationship between an individual’s mean accuracy advantage for the goal item, and their persistence, using Spearman’s correlation.

#### fMRI acquisition

The fMRI data were collected at the Oxford Centre for Human Brain Activity using a 3T Siemens scanner with a multiband accelerated echoplanar imaging sequence with the following parameters: voxel resolution 2.4 x 2.4 x 2.4 mm^3^, repetition time=1230 ms, echo time=30 ms, flip angle=60 °, field of view=240 mm, multiband acceleration factor=3, PAT factor=2, encoding direction=PA. A tilt angle of 30° was used to minimize signal drop out in the orbitofrontal cortex (Deichmann et al., 2003). Data was collected in two consecutive runs of approximately 25 minutes, where participants stayed in the scanner between runs.

#### Pre-processing and analysis structure

Data were pre-processed using FMRIB’s Software Library (FSL), using the FEAT software tool (Woolrich et al. 2001). Functional data were motion corrected using rigid body registration to the central volume (Jenkinson et al., 2001, 2002). Gaussian spatial smoothing was applied with a full-width half-maximum of 5mm, and high pass temporal filtering was applied with a cut-off of 60s. Cardiac and respiratory data were processed using FSL’s Physiological Noise Modelling (PNM) tool to model the effects of physiological noise in the MRI data (Brooks et al. 2008). Since participants completed the MRI session in two runs, parameter estimates were first estimated at the level of run (first level), then combined within individuals as Fixed Effects (second level), and finally combined across subjects using FMRIB’s Local Analysis of Mixed Effects (FLAME1+2; third level; Woolrich et al. 2004). Multiple comparisons were corrected for using a Z statistic threshold of 3.1, and a cluster probability threshold of p=0.05.

#### Univariate fMRI analyses

##### Decision-time analysis

A general linear model (GLM) was used to model BOLD activity in pre-whitened data space. Seven regressors of interest were included in the main GLM, predicting BOLD activity at the onset of the decision period (all modelled as stick functions). These regressors included whether the choice on this trial was to persist or abandon (coded as 1/-1), the full-task value of the current goal, the full-task value of the two alternatives, goal progress, goal size, and reaction time. Since goal progress is both correlated with full-task value, and our behavioural analyses shows it is an additional predictor of abandonment beyond full-task value (illustrated in Fig.2c), we disentangled the goal progress component from value in the MRI analysis. To do this, we residualised all forms of value to goal progress, and used goal progress as an independent regressor allowing us to identify where goal progress is separately tracked in the brain. In addition, since the full-task value of an option is an approximation of its ‘time to completion’, it is highly dependent on the size of the net across different blocks. To account for this, we also residualised full-task value to net size, and included net size as a separate regressor. In other words, for each value component (current goal, best alternative, worst alternative), we removed the components related to goal progress and goal size, and added these components as unique regressors. All regressors were z-scored at the level of individual runs before fitting the GLM. Supplementary fig.6a displays the final correlations between the regressors.

In addition to the parametric regressors, five types of events were included in the final GLM as main effects: onset of the decision period, onset of the block, spatial presentation of the three stimuli (substituting the spatial task), the update of the net, and the end of the block. The following confound regressors were also included in the design matrix: Six motion regressors produced during realignment, the physiological explanatory variables (processed by PNM) and motion outliers detected using FEAT’s *fsl_motion_outliers* tool.

##### Whole-brain inter-trial analysis

Given that behavioural biases accompanying goal pursuit lasted even outside of the decision period (in our spatial task), we asked whether goal-related neural activity persisted between decisions too. Of the regressors listed under *Univariate fMRI analyses,* goal progress is the one dynamic variable which can be tracked between trials (rather than depending on information presented at the decision; i.e. the offers which feed into the option values). We therefore specifically investigated whether information about goal progress was carried between trials.

To do this, we ran a whole brain analysis where we included all the same regressors listed in *Decision-time Analysis*, both time-locked to the decision onset *and* time-locked to the presentation of the first fixation (ITI 1; see Supplementary fig.1d for ITI timing during task; see Supplementary fig.6d for regressor correlation matrix). We asked whether the activity tracking goal progress was present during the inter-trial interval (see Fig.3b for results of this analysis; See Supplementary tab.2 for results relating to inter-trial goal progress activity).

#### Region of interest analyses

##### ROI selection and extraction procedure

We selected ROIs in vmPFC, ventral striatum, and ACC on the basis of value-related activity peaks at the decision time. This involved selecting peaks either for activity related to the contrast capturing the value persisting (current goal value–best alternative value; peaks in vmPFC and ventral striatum), or capturing the value of abandonment (best alternative value– current goal value; dACC), following cluster correction (Illustration of ROIs in Supplementary fig.7a,b,c; all activity peaks listed in Supplementary tab.1). Since our whole-brain analysis did not reveal any activation for the value of the third alternative in these areas, we did not include the third alternative in subsequent analyses. Regions of interest consisted of spheres with a 3 voxel radius (7.2mm^3^). In time-course analyses, activity in these spheres was up-sampled by a factor of 10, and cut into epochs which were aligned to the onset of the decision phase (see plots of activity time-courses in Supplementary fig.d,e,f).

Activity in these value-related ROIs was then used to investigate a) the modulation of value signals over the course of goal progress and b) correlations with individual differences in persistence biases. Any time courses displaying non-orthogonal contrasts are for illustration purposes only and no statistical tests were performed.

##### Baseline activity analysis

Our previous whole-brain analysis found that activity relating to goal progress was present in the inter-trial interval, with the peak of this activity located in vmPFC. Previous research has shown that baseline representations of long-term task variables influence subsequent choice behaviour through vmPFC (Vinckier et al. 2018, Abitbol et al. 2015, Lopez-Persem et al. 2016). We therefore asked whether this baseline goal-related activity at the onset of decisions was relevant for the behavioural differences in choices and attention we observed.

We quantified individuals’ baseline representation of goal progress in the vmPFC ROI (see ROI selection and extraction procedure). As in (Lopez-Persem et al. 2016, Vinckier et al. 2018), we define baseline activity as the activity present at the onset of the choice offers, before the new offers or decision itself influence the dynamics (i.e. t=0 of the time course shown in Supplementary fig.7d,e,f). We predicted vmPFC baseline activity in a model with all the identical regressors to those listed in the whole brain analysis (see *Decision Time analysis).* Then we specifically tested for a relationship between the beta-weight for goal progress (proportion of goal completed) and our behavioural measures (persistence bias and attentional goal capture).

To test the specificity of our vmPFC baseline effect we did two additional control analyses. First, we tested whether baseline representations of goal progress in the other two ROIs (ventral striatum and ACC) significantly predicted our persistence biases (Supplementary fig.8b). Second, we tested whether the effect was specifically driven by baseline rather than decision-related activity, by investigating whether goal-related activity time-locked to the decision itself predicted individual behavioural measures (Supplementary fig.8c). To quantify the decision-related activity, we multiplied the fitted beta coefficients for goal progress at each time-point by the double gamma HRF function, and summing the products to produce a coefficient for each participant (same procedure described in *Value modulation analyses*).

##### Value modulation analyses

We found an asymmetry in the use of value in behaviour, where the influence of value related to alternative goals disappeared more than the influence of value related to the current goal, over the course of goal pursuit. Therefore, we asked whether neural representations of value also changed over the course of goal pursuit.

We began by investigating whether any of our ROIs displayed decreasing impact of value-difference over the course of goal progress. As for behaviour, we predicted neural activity using the interaction between goal progress and abandonment value (value of best alternative–value of current goal). We included regressors for the main effects as well as additional regressors controlling for switch choices and reaction times (log RT). All regressors were normalised before fitting the GLM. To test for statistical significance, we multiplied the fitted beta coefficients for the interaction term (goal progress* alue) at each time-point by the double gamma HRF function (also used in the whole brain analysis) and summed the products to produce a coefficient for each participant. These were then tested against 0 using the Wilcoxon signed-rank test.

We then investigated whether a modulation in value representation was driven more by one value component over the other. We split abandonment value into its two components, namely the value of the current goal, and the value of the best alternative, and modelled their interaction with goal progress separately. As with the previous analysis, we included reaction time and abandonment trials as additional regressors, as well as including all the main effects of the interaction terms. The same Wilcoxon test described above was performed to determine whether there was a significant change across individuals in value representations for alternatives (goal progress*best alternative value) or for the current goal (goal progress*current goal value).

### Lesion Patient Study

#### Participants and experimental procedure

Twenty-six patients with brain lesions (mean age=58) and twenty-seven age-matched control participants (mean age=59) took part in the study. Of the lesion patients, one was excluded because they failed to pass the initial comprehension questions, and two were excluded because they were unable to complete the task. Of the remaining twenty-three individuals in the study, sixteen had damage within the frontal cortex and the remaining seven had damage to other areas (See Fig.5a for maps of lesion overlap). The patient population was recruited from a database of individuals who had previously visited the John Radcliffe Hospital and consented to be contacted for research studies. Data collection took place online, over a single session where the participant completed an online version of the task (hosted on Pavlovia), while the researcher remained on the telephone throughout the session. Before beginning the task, the participant received 12 trials of training, and was required to pass three comprehension questions before proceeding to the main task, which consisted of 250 trials total. The same schedule was used across all participants. The age-matched controls completed the same schedule and training procedure online, and were recruited through Prolific.co.

#### Voxel-wise lesion analysis

We began by investigating the relationship between brain damage and persistence biases independently from the fMRI study. To investigate areas causally relevant for persistence in the task, we performed a voxel-wise whole-brain analysis predicting behaviour from maps of the patients’ neural damage (Fig.5b). For each voxel, we predicted individual persistence biases using a binary regressor capturing whether the voxel was damaged in that individual:

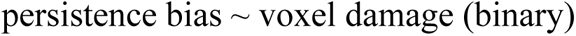

We used a threshold of *t*>2.3 where damage predicted lower persistence biases (p<0.01, one-sided test because we were interested in where damage will *reduce* persistence biases).

##### Permutation-based cluster correction

We controlled for multiple comparisons by performing cluster correction using the False Discovery Rate method (FDR; Genovese et al. 2002). Using a permutation-based approach, we asked what the maximum cluster size we would expect from our lesion dataset due to chance was, at the same significance threshold. On each permutation (total 1000 iterations), we shuffled individual persistence biases and performed the same voxel-wise regression analysis with the shuffled biases. We created a distribution of clusters found across all permutations, and defined the minimum cluster size for significance at the 95% cut-off of all clusters found by chance, resulting in a minimum cluster size of 255 voxels.

#### ROI-based lesion analysis

Next, we performed a group-wise comparison where we split lesion patients based on whether they were damaged in a region pre-defined by our fMRI study. Our fMRI study had identified a subset of areas carrying signals relating to goal-pursuit even between decisions, focussed on vmPFC. We split all patients into two groups on the basis of whether they were damaged at an ROI centred on the peak of this inter-decision fMRI activity. Following the same procedure described in ROI selection and extraction procedure, we extracted a region of interest with a 3 voxel radius (7.2mm^3^) centred on the peak of activity tracking goal progress during the inter-trial interval in our fMRI study. We then tested for a difference in persistence biases between the two groups of patients, and against the age-matched controls. We used a one-sided permutation test to test for difference in means between groups, due to the small sample sizes and non-normally distributed biases (Fig.5c, we used a one-sided test based on our hypothesis that damage to vmPFC would reduce persistence although note the difference remains significant if we were to perform a two-sided test).

#### Patient control analyses

Our voxel-wise regression analysis identified a region of vmPFC which included damaged voxels from five different patients. Our ROI-based lesion analysis independently identified four out of the five same patients when selecting on the basis of a pre-defined fMRI region. For the subsequent control analyses, we verified that the initial five patients were truly less biased to persist, rather than persisting less for other reasons (such as using a drastically different strategy, or responding more randomly). We note that if these control analyses are limited to the four patients identified in the ROI-based analysis (excluding the additional vmPFC patient identified in the voxel-wise analysis), the same conclusions hold.

First, we compared performance across groups. If vmPFC patients are truly less *biased* to persist than other patients, rather than just being more random in their switch behaviour, we should expect to see a performance enhancement. We quantified performance as the mean number of trials taken to complete a goal, where a lower value means goals were completed faster. Since all participants in the patient task completed the identical schedule, this measure is not vulnerable to schedule-specific artefacts. We then tested whether vmPFC patients performed better than patients with damage elsewhere, using a one-sided parametric test (Fig.5d; we used a one-sided test based on our hypothesis that reduced bias should improve performance but note the difference remains significant if we were to perform a two-sided test).

Second, we confirmed that behaviour among patients with damage to this region was still best explained by the same behavioural model as healthy individuals (the optimal ‘full-task model’), and not by a more simple strategy, by fitting the four behavioural models in the same way as described in *Fitting Normative Models* (Supplementary fig.9a).

Finally, we verified that the vmPFC patients were not more stochastic in their decision process. We quantified stochasticity as inverse temperature, which is the beta weight associated with the optimal value in our logistic regression predicting abandonment from optimal value. We used two-tailed permutation tests to verify there was no difference in stochasticity between the vmPFC lesion group and other patients, and between the vmPFC lesion group and age-matched controls (Supplementary fig.9b).

#### Spatial task in lesion patients

Our patient group also performed the interleaved spatial task. We quantified spatial attention bias as the accuracy advantage for the current goal item over the alternative item, as described in *Spatial task analyses.* We predicted the vmPFC group would show a lower accuracy advantage for the goal item over the alternative items in the interleaved task, since attention would not be captured by the goal.

While as predicted, the vmPFC group did not show a significant accuracy or reaction time advantage for stimuli related to the current goal (goal item accuracy advantage: mean=0.026, std=0.031, Wilcoxon signed-rank for difference against zero: *n*=5, *T*=2.0*, p*=0.188; goal item reaction time advantage: mean=-0.017, std=0.116, Wilcoxon signed-rank for difference against zero: *n*=5, *T=*5.0, *p*=0.625), we cannot interpret this result since we also did not detect goal-oriented spatial attention effects among patients with lesions elsewhere either (goal item accuracy advantage: mean=0.031, std=0.130, Wilcoxon signed-rank for difference against zero: *n*=18, *T*=80.0, *p*=0.832; goal item reaction time advantage: mean=0.041, std=0.104, Wilcoxon signed-rank for difference against zero: *n*=18, *T*=5.0, *p*=0.054). Since we were unable to detect goal-oriented attentional biases in either group, there was also no difference in attentional biases between groups (permutation test for difference in goal item accuracy advantage across groups: mean difference=0.004, *p*=0.464, *n.s.*; permutation test for difference in goal item reaction time advantage across groups: mean difference=0.058, *p*=0.297, *n.s.*).

A likely explanation for the difficulty detecting attentional biases in the patient cohort compared to the MRI cohort is simply that the fast-paced spatial attention task was too difficult for the older brain-damaged population. In general, this is reflected in accuracy: accuracy among the patient group was considerably worse compared to our MRI participants (mean error in fMRI sample: *n*=30, mean=0.210 screen units, std=0.026, mean error in patient sample: *n*=23, mean=0.316, std=0.182; permutation test for difference in means: mean difference=0.106, p=0.002; see Supplementary fig.9e for raw error in each group). In addition, unlike with the MRI cohort, this task was performed remotely with the patients, with likely variation in computer and mouse set-up and internet speed hampering the ability to detect subtle differences in responses in the spatial task. Given we could not detect goal-oriented attentional effects in the lesion patient population for the reasons discussed, we cannot determine whether lesion location affects spatial attention in this study.

