## Supplementary Materials for "VmPFC supports persistence during goal pursuit through selective attention"

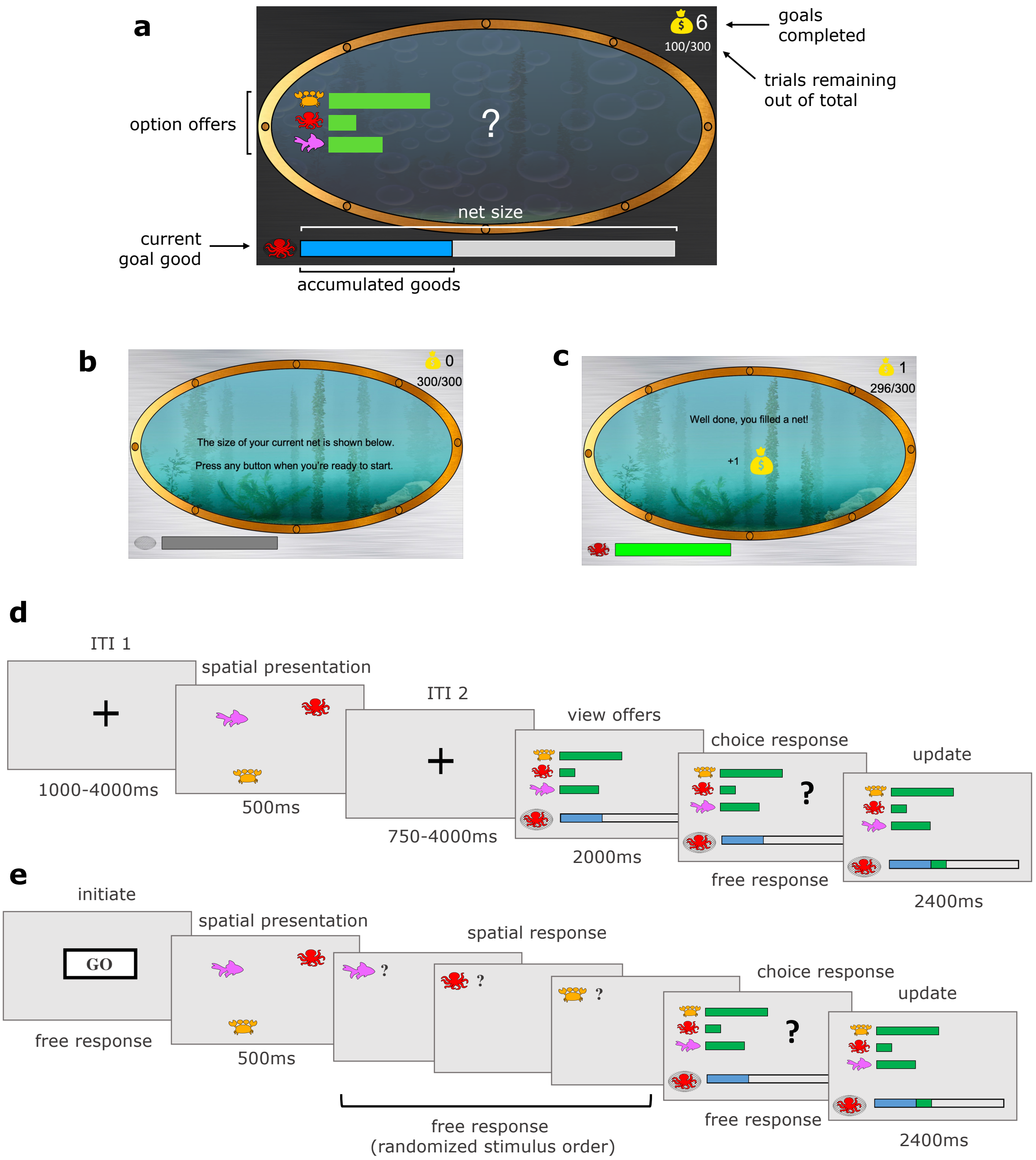


**Supplementary Figure 1. Task Design.**

**(a)** Illustration of task as presented to participants using the framing story of an 'underwater' fishing game. In addition to the features described in Fig.1a, the number of nets completed (i.e. points) was shown in the top right corner next to the money bag icon. Total nets were directly translated to the participant's bonus payment at the end of the study. The number of trials remaining in the session was shown directly underneath, as a proportion of the total trials in the session. This was to incentivise participants to make as strategic choices as possible to maximise the number of nets they could fill within the remaining trials.

**(b)** At the start of each new block, participants were presented with the size of the net they needed to fill.

**(c)** A block ended when the participant had filled the net, and a point was won.

**(d)** Task sequence inside the scanner. To keep the task visually consistent with the spatial session outside the scanner, participants passively viewed the three sea creatures flash on screen during the inter-trial interval, but were not required to report the location of the creatures. To dissociate activity related to the decision from activity related to response indication, we included a two second buffer zone once the offers were presented, before participants could make their response. In the main fMRI analyses, activity was time-locked to the onset of the decision period, shown here as 'view offers'. In the additional ITI analysis, activity was time-locked to 'ITI 1'.

**(e)** Task sequence in the spatial session (outside the scanner). Participants initiated the presentation of the creatures. After viewing the presentation of the creatures for 500ms, they were then probed on the location of the three creatures in a randomised order before being presented with the main decision task.

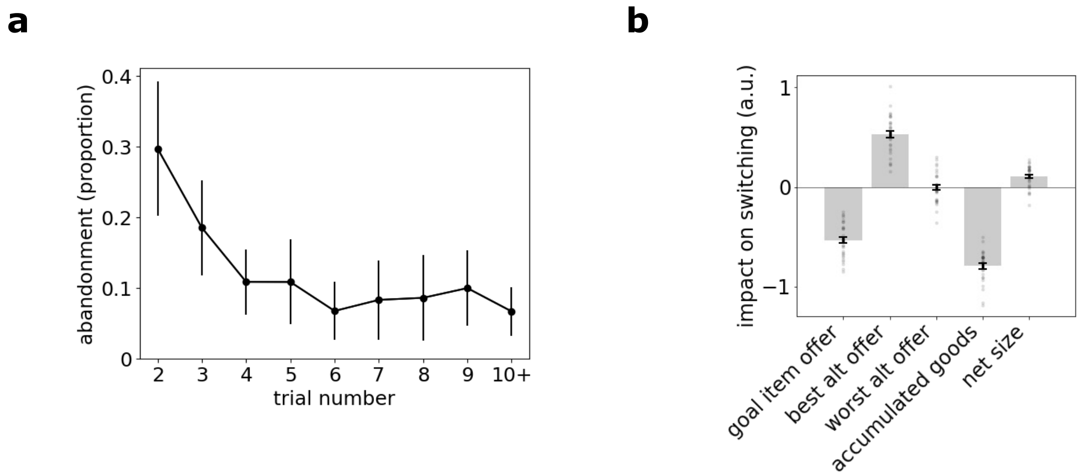


**Supplementary Figure 2. Additional behaviour**

**(a)** Proportion of abandonment choices as a function of trial number. Error bars show standard deviation across participants.

**(b)** Results of a simple logistic regression analysis, showing participants were sensitive to the key elements of the task: they were more likely to switch when the offers associated with alternative goods were high ($t(29)=14.74$, $p<0.001$), and less likely to switch when the offers for their current good were high ($t(29)=-16.84$, $p<0.001$), or after having accumulated many goods in their net ($t(29)=-27.67$, $p<0.001$). We also found that people were more likely to abandon a goal when the size of the target net was larger ($t(29)=5.35$, $p<0.001$). We found no effect of the second-best alternative on abandonment decisions.

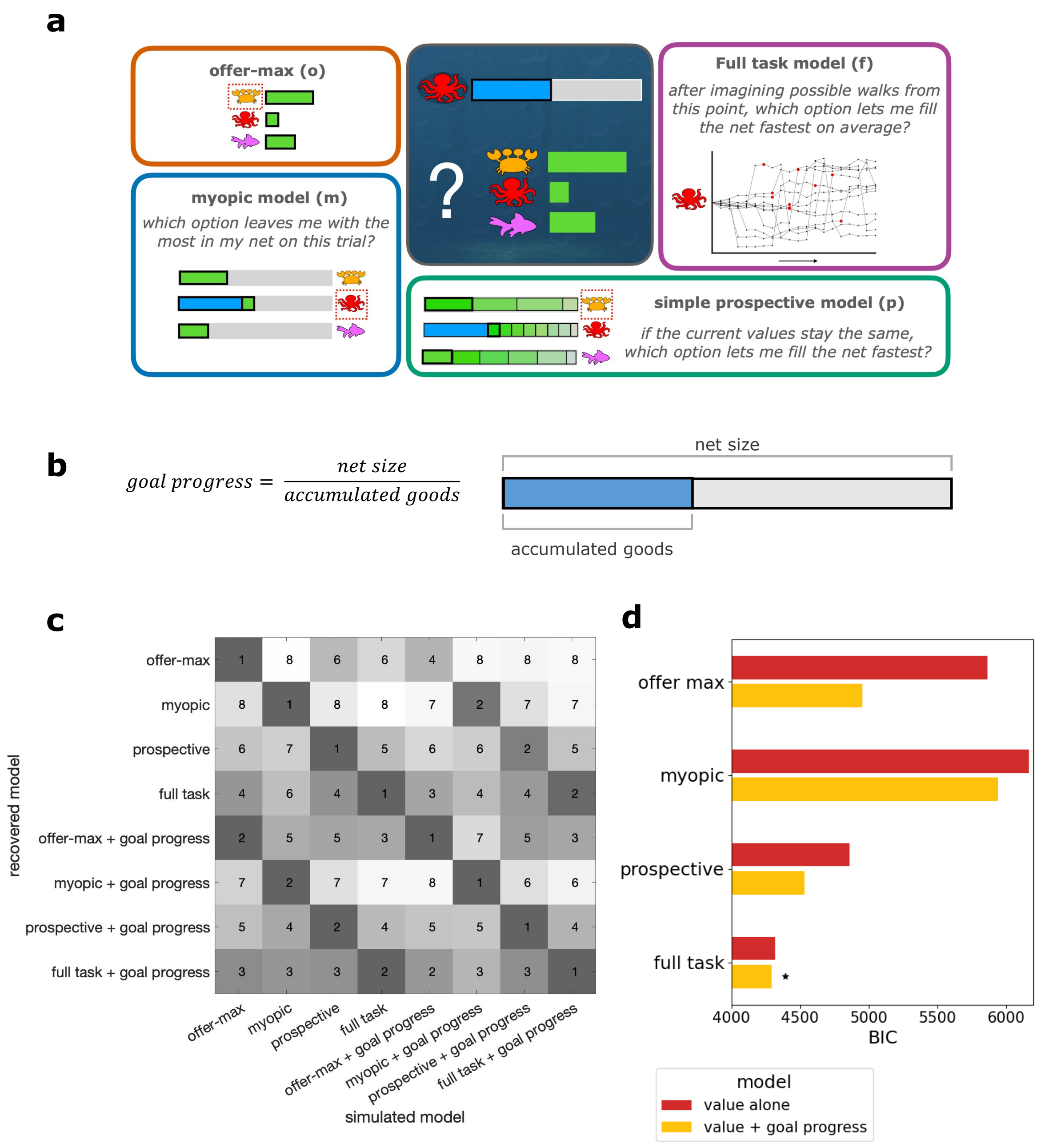


**Supplementary Figure 3. Behavioural modelling**

**(a)** Graphic of the four behavioural models as described in 'Behavioural Models' in methods.

**(b)** Graphic of the 'goal progress' regressor.

**(c)** Confusion matrix resulting from model recovery procedure. Each column corresponds to a model used to simulate the dataset. Each row corresponds to the model used to recover the dataset. Within a column, shading corresponds to the BIC of each competing model relative to the winning model. Lower BICs corresponding to better fits are displayed in darker shades. Numbers indicate the rank of the model in the model comparison per column (where 1 is the winning model, and 8 is the worst fitting model). In all cases, simulated behaviour is best fit by the true generative model.

**(d)** Behavioural information criteria (BICs) for the models fitted to participant behaviour. Each model fitted corresponds to a model recovered in (c). Dark orange depicts logistic regression models using model value alone. Light orange depicts logistic regression models with goal progress added as an additional regressor.

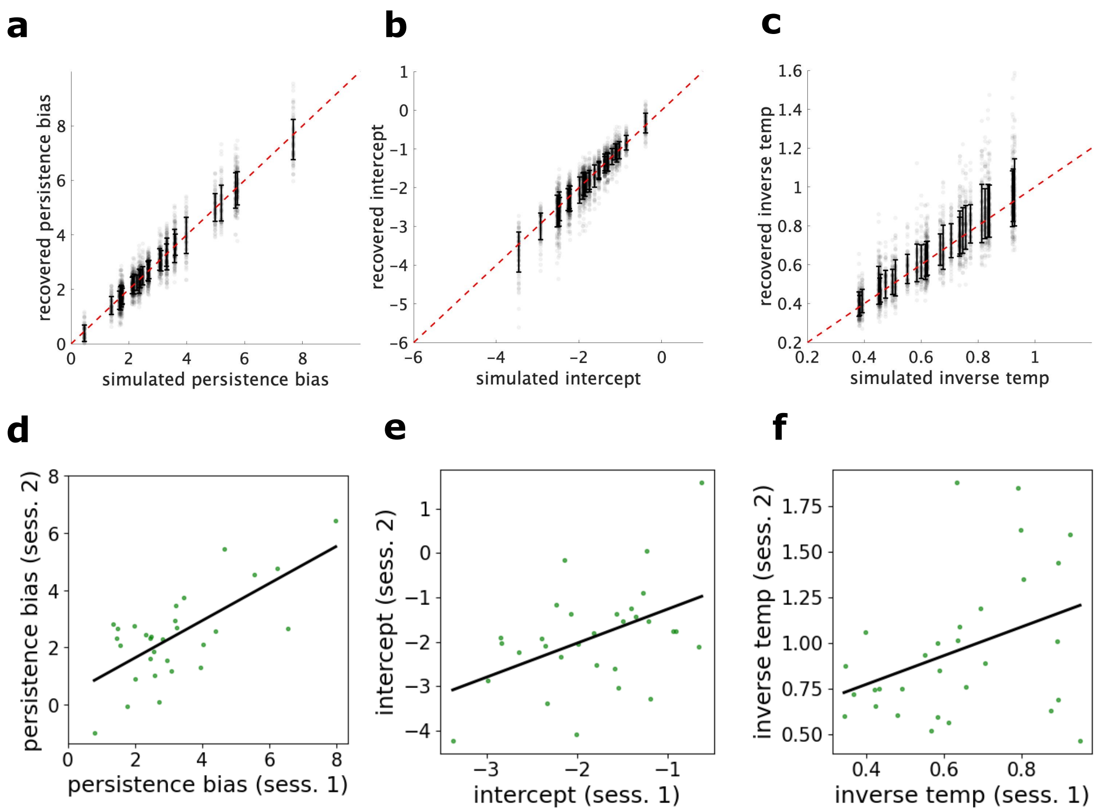


**Supplementary Figure 4. Parameter recoveries and test-retest reliability.**

**(a, b, c) Parameter recoveries**. Empirical parameters from decision data aggregated across both sessions were used to simulate behaviour (100 iterations per participant yielding 300 simulations total). Parameters were recovered for each simulation by fitting a logistic regression using the same procedure used for the empirical data. **(a)** An individual's persistence bias was defined as their 'indifference point' to abandonment when predicting abandonment choices using the full-model value of abandonment. This is equal to the negative product of temperature and intercept from the logistic regression (-intercept*temperature; see 'Persistence bias' section in Methods). Recovered persistence biases correlated with the simulated biases with a Pearson's correlation of 0.96 ($p<0.001$). **(b)** Recovery of the intercept parameter: Recovered against simulated parameters from the logistic regression. The simulated intercepts can be recovered with a Pearson's correlation of 0.92 ($p<0.001$). **(c)** Recovery of the inverse temperature parameter: Recovered against simulated inverse temperature. The simulated inverse temperature can be recovered with a Pearson's correlation of 0.86 ($p<0.001$).

**(d, e, f) Test-retest reliability of parameters across the two sessions.** Parameters were separately fitted to the decision task inside the scanner ('session 1', see supplementary fig.1d for task) and to the decision task outside the scanner ('session 2', see supplementary fig.1e for task). All parameters show significant test-retest reliability. **(d)** Test-retest reliability for persistence biases across the two sessions (Pearson's $r$=0.70, $p$<0.001). Persistence biases are derived from the intercept and inverse temperature shown in e and f. **(e)** Test-retest reliability for the intercept alone across the two sessions (Pearson's $r$=0.46, $p$=0.010) **(f)** Test-retest reliability of inverse temperature across the two sessions (Pearson's $r$=0.38, $p$=0.040).

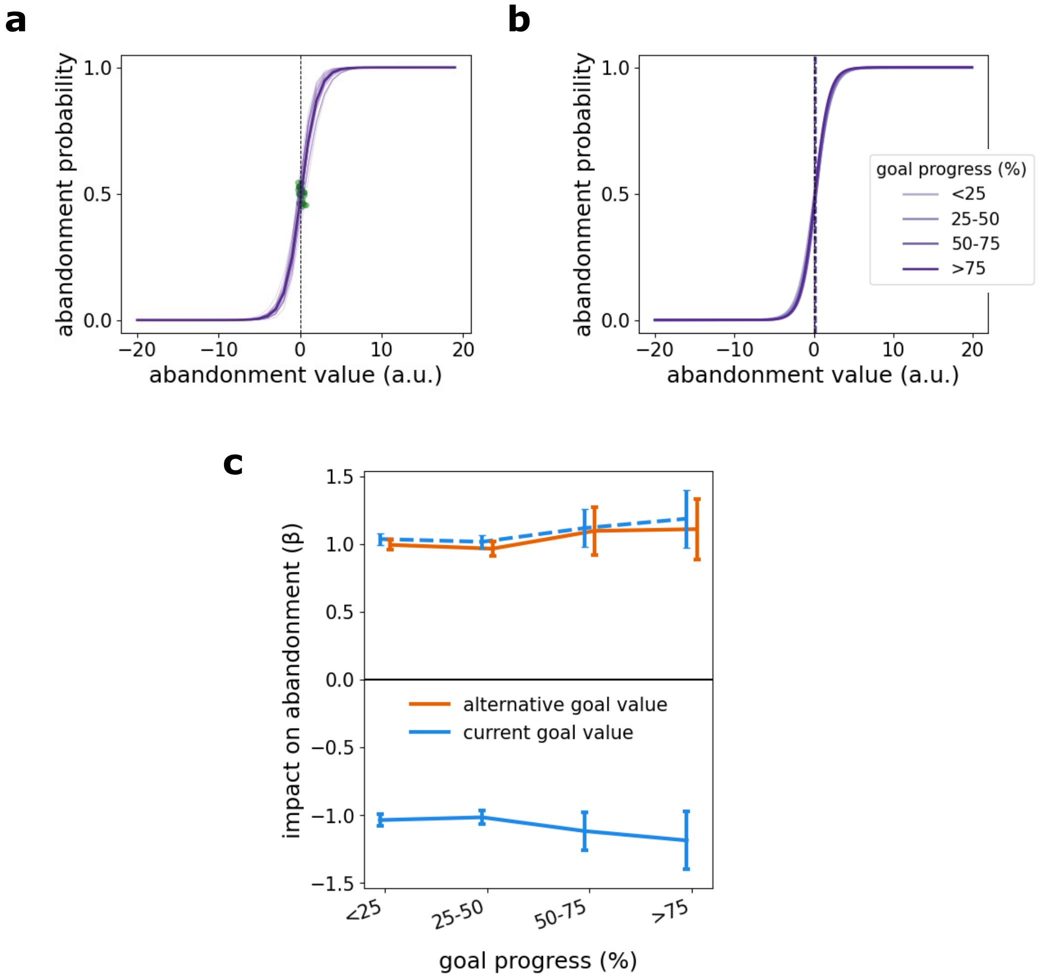


**Supplementary Figure 5. Simulated normative behaviour.**
Behavioural analyses on simulated data from the normative full-task model, demonstrating the empirical behavioural patterns are not an artefact of the testing schedules. Here, we fit the normative full-task model to 30 simulated data-sets in which each simulation corresponded to one of the the 30 participant schedules.

**(a)** Individual fits to simulated data sets show persistence biases of zero, demonstrating that the empirically found biases are not an inherent feature of the schedules used. Compare to fig.2b showing the range of participant specific persistence biases.

**(b)** Quartile fits demonstrating that we accurately recover no difference in persistence biases across quartiles when simulating with the normative model. Compare to fig.2c showing that participants are more biased to persist as goal progress increases.

**(c)** Effects of temptation and frustration on abandonment choices. In the normative model simulations, we accurately recover no difference in how these two abandonment causes are weighted. Compare to participant behaviour in fig.2d showing diverging slopes corresponding to the impact of frustration vs temptation across goal progress.

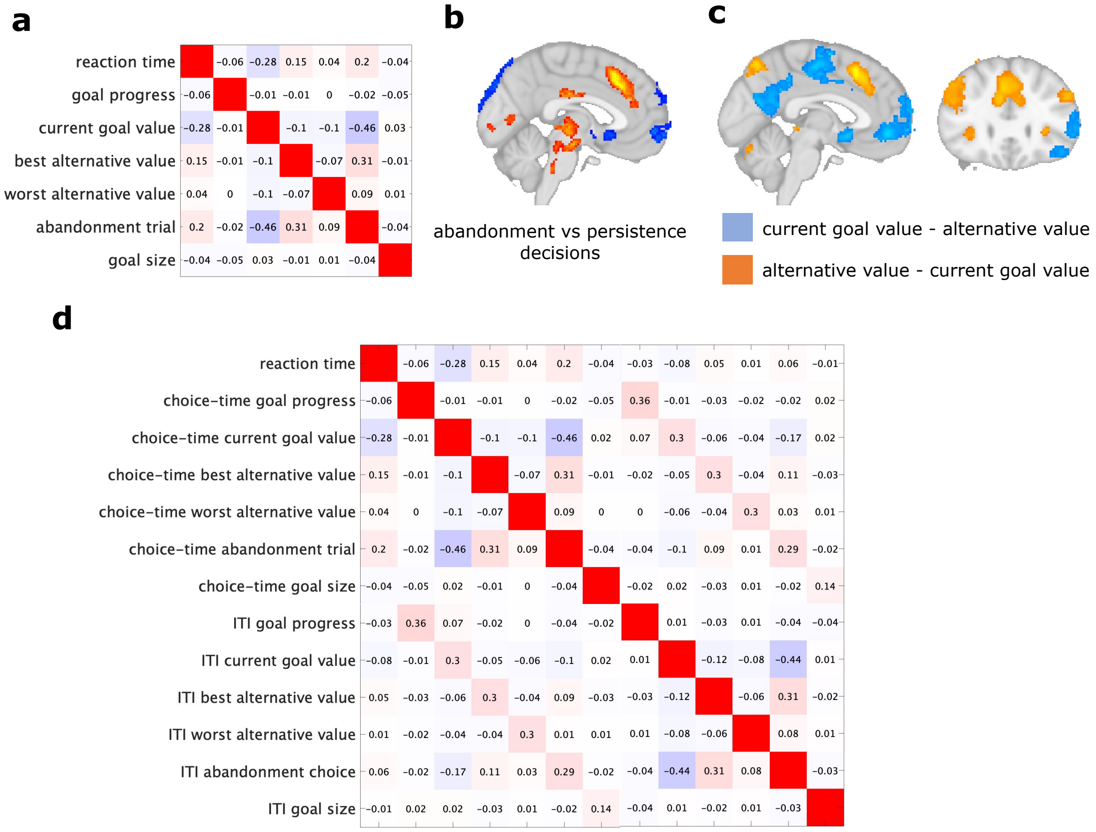


**Supplementary Figure 6. Whole-brain analyses.**

(**a**) Correlation matrix (convolved with the hemodynamic response function) of the regressors included in the initial whole-brain analysis, alongside event regressors. The regressors were time-locked to the onset of the choice (i.e. 'view offers' in Fig.1d). Value refers to value from the full-task model, the normative model which described behaviour best. Value was orthogonalised to goal progress to separate these components each shown to affect behaviour (see Methods).

**(b)** Activity related to abandonment (red) vs persistence (blue) decisions. In other words, the positive and negative activity related to the 'abandonment trial' regressor in (a), where trials were coded as 1 for abandonment and -1 for persistence

**(c)** Activity related to the value contrasts. Blue depicts areas where activity increases when the full-model value of persisting with the goal is greater than the full-model value of switching to the best alternative. Orange shows the inverse (where activity increases when the full-model value of switching to the best alternative is greater than the full-model value of persisting with the current goal). The peaks of these value contrasts were used to define the regions of interest in vmPFC, ventral striatum, and ACC, as shown in Supplementary fig. 7.

**(d)** Correlation matrix (convolved with the hemodynamic response function) of the regressors included in the second ITI whole-brain analysis. This analysis included all the same regressors as the main whole-brain analysis, this time modelling BOLD activity both at the choice onset, and in addition at the first inter-trial fixation cross (i.e. 'ITI 1' in Fig.1d).

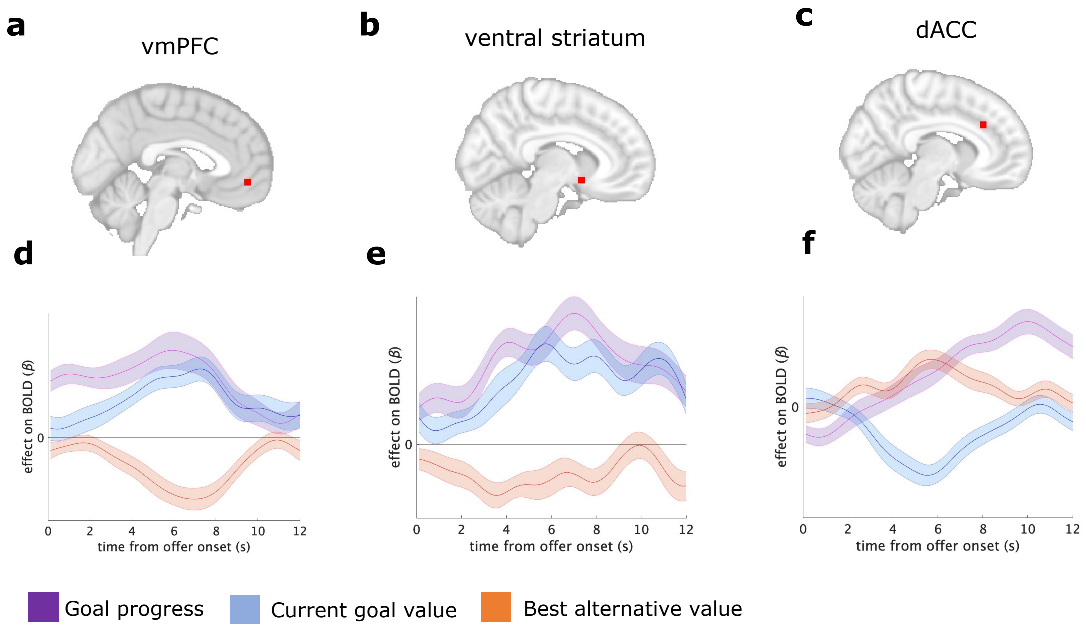


**Supplementary Figure 7. Regions of interest and time courses.**

**(a, b, c)** We extracted regions of interest based on the peaks of value-related activity in our fMRI study (see '*ROI selection and extraction procedure'* in Methods). These consisted of the peaks of activity for the contrast of goal value–alternative value in the case of vmPFC [-2,48,-8] and ventral striatum [8,8,-10], and the largest sub-peak of activity in the dACC for the alternative value–goal value regressors [8,28,30]. All peaks are shown in Supplementary Table 1 (ROI peaks starred).

**(d, e, f)** Time course analyses depicting the t-statistics for the regressors of goal progress (purple), current goal value (blue), and best alternative value (orange) in the three regions of interest (for illustration). Time 0 seconds corresponds to the onset of the choice (i.e. 'view offers' in Supplementary fig.1d). The GLM used in the time-course analysis contained identical regressors to the whole-brain GLM described in *Univariate fMRI analyses* in Methods, and shown in the correlation matrix in Supplementary fig.6a. Error bars show standard error of the mean across participants. Note the pre-decision modulation of activity by goal progress ($t$=0) in the vmPFC predicted individual differences in attention and decision (Fig.3d,e).

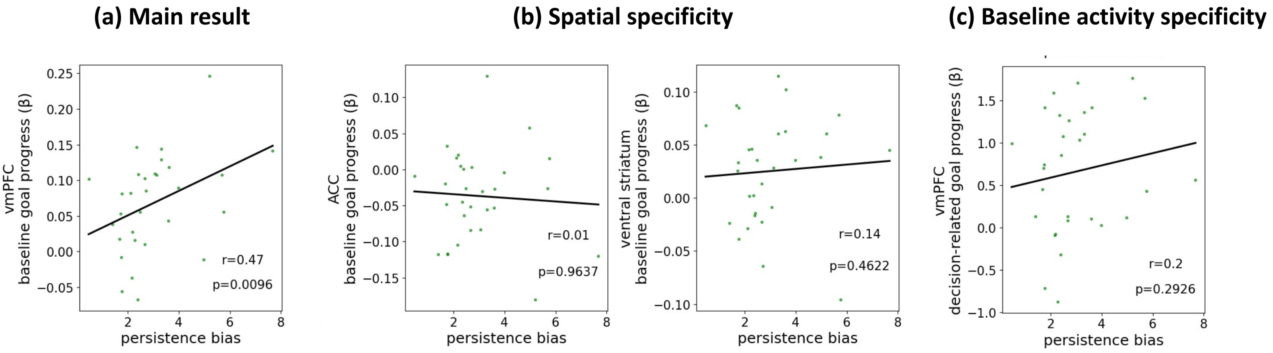


**Supplementary Figure 8. Control analyses for the relationship between baseline vmPFC activity and persistence bias.**

Statistics shown in each plot report the spearman's correlation (effect and significance) between the neural regressor, and the persistence bias fitted to data aggregated across both sessions. In all plots, green dots show individual participant biases plotted against the relevant neural regressor beta weight.

**(a)** Main effect from manuscript, showing baseline vmPFC tracking of goal progress predicts individual differences in persistence bias (where baseline representation of goal progress is defined as the beta weight for the relationship between vmPFC BOLD activity and goal progress at decision onset). Our decision to examine baseline vmPFC activity was based on hypotheses from previous literature showing baseline vmPFC activity carries subjective biases in decision-making, but here we present various controls.

**(b)** Control analysis showing spatial specificity of the effect in (a) to vmPFC: Baseline representation of goal progress in the ACC and striatal regions of interest does not predict individual differences in persistence biases.

**(c)** Decision-related representations of goal progress in vmPFC does not predict individual difference in persistence bias. Here, we capture decision-related representation of goal progress by multiplying the fitted beta coefficients for goal progress at each time-point from choice onset by the double gamma HRF function, and summing the products to produce a decision related component for each participant.

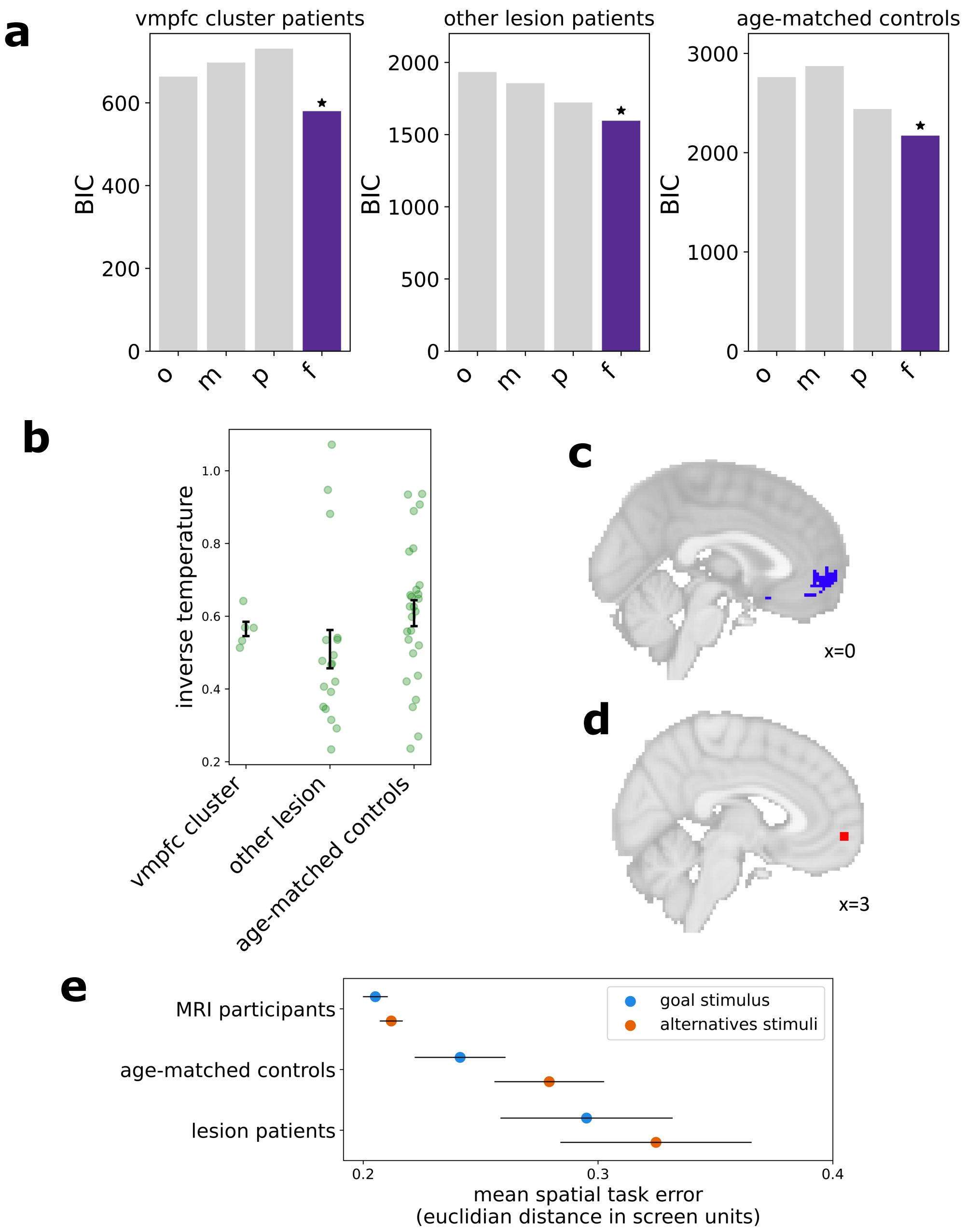


**Supplementary Figure 9. Lesion patient study.**

**(a)** Results from a post-hoc control analysis where we fit the four normative models to behaviour in the three groups (left: patients with damage within the vmPFC cluster identified from the voxel-wise analysis; middle: patients with damage external to the vmPFC cluster; right: age-matched healthy control participants). In each group, we fit the models using a mixed effects regression model predicting abandonment choices (same method as for fMRI participants, described in Methods). Our results show the full-task model remains the best fit to behaviour in all three groups.

**(b)** Post-hoc analysis comparing inverse temperature (beta weight for sampler value in the logistic regression) across the three groups described in (**a**). Patients with damage within the vmPFC cluster identified in the voxel-wise analysis are not simply more stochastic since they show no difference in inverse temperature to the age-matched controls. Note the inverse temperature parameter has high parameter recoverability (Supplementary fig.4c).

**(c)** Voxel-wise map of t-statistics where lesion damage predicts reduced persistence biases (same as shown in Fig.5b thresholded at a higher significance level for illustration, $t>2.6$).

**(d)** Region of interest from fMRI study, taken at the peak of activity tracking goal progress between decisions, at [4,58,-6] (see Supplementary table 2). Patients are split into those with damage inside this ROI and those with damage external to this ROI in Fig.5c.

**(e)** Response errors in the spatial task for each group, where error is mean Euclidian distance between participant clicks and true item locations, in normalised screen units. Orange shows average error for the current goal item, blue shows average error for the alternative items. Error bars depict standard error of the mean. While the spatial bias effect is in the right direction, the patient group do not show a significant accuracy advantage for the current goal item relative to alternative goal items.

**Supplementary Table 1**. **Peaks of activity from cluster-corrected whole-brain fMRI analysis locked to decision onset.** Only regressors of interest are reported here although see correlation matrix in Supplementary fig.6a for all regressors included in GLM. Related to Fig.3a and Fig.4a of the main manuscript, as well as Supplementary fig.6a,b,c.

| Contrast | Region | Peak coordinates (x,y,z in mm MNI space) | Z Value |
|---|---|---|---|
| **Persistence value**: current goal value–best alternative. value | Ventromedial prefrontal cortex (area 11m) | -2, 48, -8* | 5.2 |
| | Ventral striatum | 8, 8, -10* | 5.16 |
| | Lateral frontal pole | -38, 30, -16 | 5.26 |
| | Supplementary motor cortex | 0, -6, 58 | 5.8 |
| | Parietal operculum cortex | Left:- 58, -38, 26; Right: 54, -28, 24 | Left: 5.98; Right: 5.62 |
| | Intracalcarine cortex | 8, -84, 4 | 6.94 |
| | Precuneous cortex | -6, -54, 12 | 5.55 |
| | Superior parietal lobule | 28, -40, 64 | 5.04 |
| | Parahippocampal gyrus | -24, -36, -14 | 4.8 |
| | Precentral gyrus | Left: -60, 0, 30 ; Right: 42, -18, 62; | Left: 4.69; Right: 4.6 |
| | Cerebellum area VIIa | 14, -64, -46 | 4.36 |
| | Cerebral crus | 26, -80, -32 | 4.56 |
| **Abandonment value**: best alternative value–current goal value | Dorsolateral prefrontal cortex | Left: -46, 44, 16; Right: 48, 36, 26 | Left: 5.74; Right: 6.27 |
| | preSMA extending into dorsal ACC | -4, 20, 46 dACC subpeak: 8, 28, 30* | 6.27 dACC subpeak: 5.4 |
| | Insular cortex | Left: -32, 20, 4; Right: 36, 18, 6 | Left: 5.77; Right: 5.49 |
| | Lateral frontal pole | 42, 44, 0 | 4.41 |
| | Inferior frontal gyrus / precentral gyrus | -44, 8, -32 | 5.74; |
| | Supramarginal gyrus | 50, -48, 48 | 6.21 |
| | Superior frontal gyrus | -26, 2, 62 | 4.76 |
| | Intracalcarine cortex | -8, -74, 8 | 6.03 |
| | Cerebellum area VI | -8, -74, -22 | 5.53 |
| | Cerebellum area VIIb | -34, -68, -56 | 4.93 |
| **Persistence choice:** Persistence trials–abandonment trials | Area 11m / Frontal medial pole | 0, 64, -8 | 4.95 |
| | Area 25 | 0, 20, -8 | 4.45 |
| | Orbitofrontal cortex (area 47) | -42, 28, -10 | 4.48 |
| | Occipital Pole | Left: -22,-104, -2; Right: 4, -88, 36 | Left: 4.3; Right: 4.6 |

| | | | |
|---|---|---|---|
| | Lingual gyrus | 10, -90, -18 | 4.49 |
| | Superior parietal lobule | -22, -46, 62 | 4.46 |
| | Central opercular cortex | Left: -56, 6, 0; Right: 62, 2, 8 | Left: 4.85; Right: 5.54 |
| | Precentral gyrus | 26, -22, 74 | 4.62 |
| | Lateral ventricle | Left: -20, -48, 16; Right: 36, -42, 4 | Left: 4.45; Right: 4.63 |
| | Cerebral crus | 18, -80, -40 | 3.95 |
| | Inferior temporal gyrus | 54, -58, -24 | 4.33 |
| | Lateral occipital cortex | -48, -76, 34 | 4.15 |
| **Abandonment choice:** Abandonment choice–persistence choice | Caudate | -12, 16, 4 | 5.57 |
| | preSMA extending into dorsal ACC | 6, 20, 44 | Right: 5.69 |
| | Dorsolateral frontal pole | -42, 42, 22 | 4.4 |
| | Lateral frontal pole (left) | -22, 62, 2 | 4.52 |
| | Posterior cingulate gyrus | -4, -20, 30 | 4.83 |
| | Insular cortex extending into frontal operculum cortex | Left: -32, 18, -6; Right: 36, 24, 4 | Left: 7.04; Right: 6.64 |
| | Occipital pole | -12, -94, -4 | 5.58 |
| | Superior parietal lobule | -38, -52, 50 | 4.41 |
| | Occipital fusiform gyrus | 24, -74, -4 | 4.81 |
| | Inferior frontal gyrus / precentral gyrus | -46, 2, 40 | 5.38 |
| | Superior frontal gyrus | -22, -2, 58 | 4.61 |
| | Lateral occipital cortex | -26, -84, 18 | 5.14 |
| **Goal progress** (i.e. proportion of net completed) | Non-region specific cluster (74958 voxels), encompassing areas stretching from lateral occipital cortex, temporal gyrus, insular cortex, striatum, cingulate cortex, and medial prefrontal areas | 10, -92, 6 | 8.1 |
| | Middle / superior frontal gyrus | -28, 34, 48 | 5.38 |
| | Brain stem | 2, -30, -42 | 4.6 |
| **Negative goal progress** (i.e. activity related to proportion of net *remaining* to be filled) | Lateral occipital cortex | -42, -64, 50 | 6.39 |
| Family-wise error cluster corrected, z > 2.3, p < 0.05<br>*Coordinates used for fMRI region of interest analyses | | | |

**Supplementary Table 2**. **Peaks of activity from cluster corrected whole-brain analysis time-locked to the inter-trial fixation cross.** Related to Figure 3b of the main manuscript. Only regressors of interest are reported here (i.e. inter-trial representations of goal progress), although see correlation matrix in Supplementary fig.6d for all regressors included in GLM.

| Contrast | Region | Peak coordinates (x,y,z in mm MNI space) | Z Value |
|---|---|---|---|
| **Inter-trial goal progress** (i.e. goal progress time-locked to the ITI fixation cross) | Frontal medial pole (Area 10 stretching to area 11 and area 14) | 4, 58, -6* | 5.17 |
| | Right hippocampus | 26, -20, -14 | 4.69 |
| | Temporal pole | Left: -40, 22, -30; Right: 48, 18, -30 | Left: 4.34; Right: 4.47 |
| | Area 8m | -20, 36, 48 | 4.73 |
| | Precuneus cortex stretching to posterior cingulate gyrus | -6, -56, 22 | 4.48 |
| | Middle temporal gyrus | -64, -12, -14 | 4.46 |
| | Caudate | 14, 20, 6 | 4.08 |
| | Postcentral gyrus | 36, -24, 56 and second cluster at 54, -22, 54 | 4.45, 4.58 |
| Family-wise error cluster corrected, $z > 2.3$, $p < 0.05$<br>*Coordinates used for lesion patient study region of interest | | | |
